# Synergy between tubulin code and MAPs enables selective kinesin motility

**DOI:** 10.64898/2026.11.23.767929

**Authors:** Ilia Zhernov, Kassandra Ori-McKenney, Agnieszka Szyk, Antonina Roll-Mecak

**Author notes:** Corresponding author: Antonina Roll-Mecak, Cell Biology and Biophysics Unit, Porter Neuroscience Research Center, National Institutes of Health, Building 35, Room 3B-203, 35 Convent Drive, MSC 3700, Bethesda, MD 20892-3700.

## Abstract

Neuronal motors must navigate a crowded microtubule surface decorated with microtubule-associated proteins (MAPs) and intrinsically disordered tubulin tails bearing extensive glutamylation. Many MAPs, including tau, linked to Alzheimer’s disease and frontotemporal dementia, form condensates on microtubules that block motors, yet how cells regulate access through these barriers is not well understood. Combining *in vitro* reconstitution and single-molecule imaging, we show that tubulin glutamylation enables processive motility of the anterograde motor kinesin-3 KIF1A through tau condensates which are otherwise impenetrable. Glutamylation drives a monotonic increase in tau diffusivity within condensates, altering their material properties to facilitate motor passage. This mechanism is motor-specific: kinesin-1 KIF5B cannot traverse tau condensates, even with its activator MAP7, while MAP9 synergizes with glutamylation to potently enhance KIF1A motility to near tau-free levels. Together, these findings establish that track-specific transport in the crowded neuronal milieu is an emergent property arising from the interplay between motor-specific properties, MAP composition, and the tubulin code.

## INTRODUCTION

Kinesin and dynein motors drive the organized transport of cellular cargo on microtubules, cylindrical polymers of αβ-tubulin heterodimers, a process that is essential to the viability of all eukaryotic cells. Nowhere is this transport more critical and demanding than in neurons, where long-range communication between distant compartments depends on the fidelity and efficiency of motor-driven delivery. To support this system, microtubule tracks in neuronal processes are stabilized and organized into densely packed bundles by an abundance of MAPs^1^, many of which share overlapping binding sites with kinesin and dynein motor domains. Among these, tau stands out because of its abundance and disease relevance – linked to Alzheimer’s disease, frontotemporal dementia and a spectrum of neurological pathologies^2–6^. Tau is present in neurons at micromolar concentrations^7–10^, nearly two orders of magnitude above its binding affinity for the microtubule^11–13^, and is particularly enriched in the axon where it plays an important role in axonal outgrowth^14^. *In vitro*, tau forms microtubule-associated condensates, stabilized by multivalent interactions mediated through its large intrinsically disordered regions, present both at its N and C-terminal regions^11,15–17^. These condensates act as barriers to both kinesin^11,15–22^ and dynein motors^15,18–21,23^, raising a fundamental question: how do neurons reconcile the inhibitory properties of tau with the need for high-volume motor traffic along the same tracks?

Molecular motors navigating this landscape also encounter a brush of intrinsically disordered C-terminal tails of tubulin, which project from the microtubule surface and are densely modified with chemically diverse posttranslational modifications. Together these modifications constitute a “tubulin code” that regulates the recruitment and activity of a wide range of molecular effectors^24^. For example, dynein/dynactin preferentially initiates retrograde transport on tyrosinated microtubules^25,26^, kinesin KIF13B motility is stimulated by microtubule tyrosination ^27^, severing enzymes selectively remodel glutamylated microtubules^28–31^, ER-resident proteins interact differentially with glutamylated microtubules to control ER distribution in response to cellular stress^32^, and microtubule plus-end binding proteins synergize selectively on tyrosinated microtubules to enhance microtubule dynamicity^33^. In neurons, tubulin posttranslational modifications are especially abundant and are organized in nm-scale patterns in axons and dendrites^34,35^, a spatial precision that underscores tight regulatory programs and the sophistication of tubulin-code based regulation.

Among neuronal tubulin posttranslational modifications, glutamylation - the addition of variable numbers of glutamates to internal positions in the α-and β-tubulin C-terminal tails - is highly abundant in neuronal processes^36,37^ and is the most prevalent modification in the brain. Introduced by multiple enzymes of the TTLL (tubulin tyrosine ligase-like) family, glutamylation accumulates monotonically during neurogenesis^38,39^ and its dysregulation is both a driver of neurodegeneration^24,40,41^ and a hallmark of nervous system disorders^42–45^. Consistent with this, defects in glutamylation homeostasis impair trafficking of diverse cargos such as mitochondria, lysosomes, synaptic vesicles and intraflagellar particles^42,43,46,47^. The glutamate chains are added to the intrinsically disordered tubulin tails, thus amplifying the electronegativity of the tubulin tail brush that is critical for the binding of tau and most neuronal MAPs^48–50^. Thus, the neuronal microtubule surface presents a composite, disordered environment: an electronegative tubulin tail brush interwoven with a mesh of MAP polypeptide chains which motors need to navigate to support high-volume and track-specific cargo traffic.

Despite the established importance of MAPs and tubulin posttranslational modifications as individual regulators of motor activity, how these two layers interact combinatorially to control motor behavior remains unknown. Glutamylation increases tau’s microtubule binding affinity^13^; however, the significance of this observation is unclear since tau is present in neurons at concentrations nearly two orders of magnitude in excess of its *K_D_*. Yet, motors must navigate this tau-dense environment efficiently. How this is achieved is a fundamental and open question. A key obstacle is that simplified *in vitro* reconstitution systems - while invaluable for dissecting individual regulatory inputs - often fail to recapitulate motor track selectivity observed in cells, pointing to emergent regulatory mechanisms that only manifest in more complex molecular environments. Here we address this gap directly, by developing an *in vitro* reconstitution platform that combines differentially glutamylated microtubules, tau condensates and auxiliary MAPs with single-molecule imaging, enabling us to interrogate for the first time the combinatorial effects of tubulin glutamylation and MAPs on the two major anterograde kinesin motors in neurons, KIF1A and KIF5B.

Using this system, we demonstrate that glutamylation alters the material properties of tau condensates on microtubules, converting them from impassible barriers into permissive environments for processive motility for KIF1A but not KIF5B - revealing an unanticipated mechanism of motor selectivity encoded at the level of the microtubule surface. We further show that MAP9 acts as a potent positive modulator of KIF1A, dramatically enhancing its motility through tau condensates in a glutamylation-dependent manner, whereas MAP7, a positive regulator of KIF5B, fails to support KIF5B motility even in the presence of glutamylation. Together, these findings establish that processive, selective transport through MAP-decorated microtubules is an emergent property arising at the intersection of motor-intrinsic properties, MAPs and tubulin posttranslational modifications, a mechanistic complexity that is accessible only through increasingly sophisticated *in vitro* reconstitution systems that capture the molecular context of the microtubule surface to reveal the principles governing cargo transport.

## RESULTS

### Glutamylation enables motor-selective transit through tau condensates

Previous work using microtubules assembled from brain tubulin, which has a heterogenous mixtures of posttranslational modifications, showed that tau binds microtubules with high affinity ^11–13^ and forms “condensates” or “envelopes”^15,16^. Condensate formation requires the intrinsically disordered tubulin C-terminal tails^15^, as well as the tau projection domains, which are involved in tau-tau homotypic interactions^15^. At low concentrations (nM), tau forms condensate “islands” – regions of high-density, immobile tau, interspersed with regions of low tau density where tau molecules are diffusive^15,16^. Tau is present at μM concentrations in neurons^7–10^, compatible with the formation of condensates as reported previously^14^. *In vitro*, tau condensates block microtubule binding by kinesin-1 KIF5B^16^, kinesin-3 KIF1A^11^ and severing enzymes^13,16^, an inhibitory effect which is consistent with the overlapping binding sites of KIF1A and KIF5B with tau on the microtubule surface (Fig. 1a). Given that neuronal microtubules are highly glutamylated and support a high volume of traffic driven by KIF1A (*inter alia*, responsible for synaptic vesicle transport), we investigated the effects of glutamylation on tau condensate formation, and in turn KIF1A motility through these condensates.

**Fig. 1.**
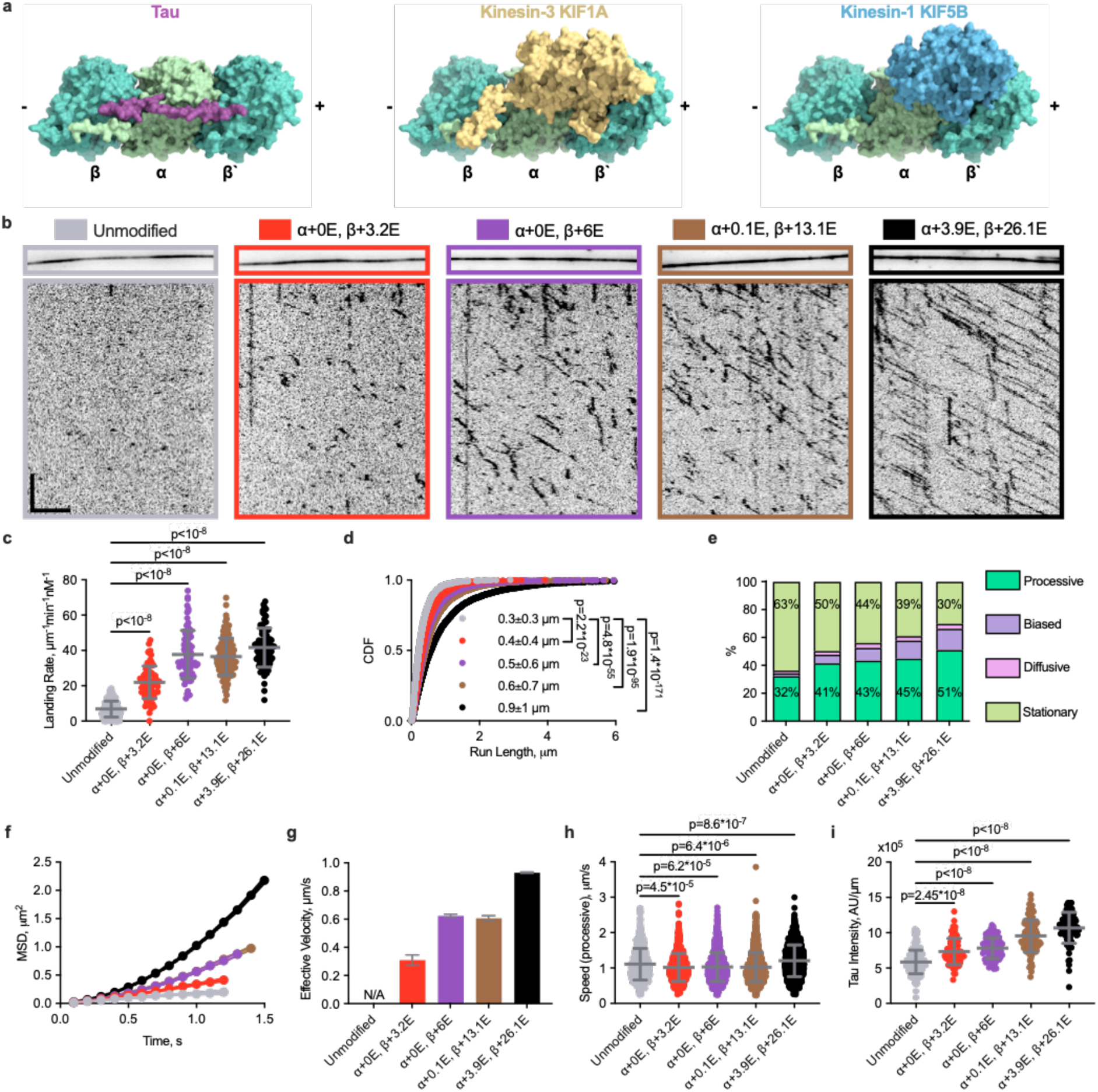
Glutamylation on β-tubulin enables kinesin-3 KIF1A processive motility through tau condensates. **a**, Cryo-EM structures of the tau R2 microtubule binding repeat (PDB 6CVN), KIF1A (PDB 8UTN), and KIF5B (PDB 8RHB) bound to a section of a microtubule protofilament. α- and β-tubulin protomers labeled in black; (-) and (+) denote the minus and plus ends of the microtubule, respectively. **b**, Kymographs of KIF1A-mScarlet motility through tau condensates on unmodified microtubules and microtubules with increasing β-tubulin glutamylation levels. 20 nM sfGFP-tau. Scale bars: horizontal, 5 μm; vertical, 5 s. **c**, KIF1A landing rate, p-values determined by one-way ANOVA. **d**, Cumulative Distribution Function (CDF) of KIF1A run lengths; mean values ± S.D. shown next to graph; n = 2174, 2366, 5052, 5417, and 4072 molecules for unmodified, <n_E_>_β_ ∼ 3.2, 6, 13.1 and 26.1, respectively: p-values determined by Kolmogorov-Smirnov test. **e**, Classification of KIF1A tracks. Classification is based on number of backsteps (N_b_) and total displacement (Δx). Processive, N_b_ < 3 and Δx > 0.3 μm; biased N_b_ > 3 and Δx > 0.5 μm; diffusive, N_b_ >3 and Δx <0.5 μm; stationary, Δx <0.3 μm. **f**, Mean square displacement (MSD) of KIF1A as a function of time; error bars, S.D., too small to be visible. See Methods. **g**, Effective velocity extracted from MSD fits in (**f**). **h**, Speed of KIF1A in processive runs, n = 759, 1055, 2376, 2681, 2317 molecules, for unmodified, <n_E_>_β_ ∼ 3.2, 6, 13.1, and 26.1, respectively. **i**, Intensity of sfGFP-tau in condensates, n = 224, 80, 85, 114, and 83 microtubules for unmodified, <n_E_>_β_ ∼ 3.2, 6, 13.1, and 26.1, respectively. 12 independent experiments for unmodified and 3 for <n_E_>_β_ ∼ 3.2, 6, 13.1 and 26.1 microtubule conditions. Average number of glutamates added to α-and β-tubulin, α + <n_E_>, β + <n_E_> throughout. Data points colored according to glutamylation levels, defined in (**b**). For microtubule mass spectra see Extended Data Fig. 1. For KIF1A motility through tau condensates, see Supplementary Video 1.

First, to investigate the relationship between tau condensate formation and tubulin glutamylation, we assembled microtubules from purified unmodified human tubulin, and then controllably modified them with the glutamylase tubulin tyrosine ligase-like 7 (TTLL7). Glutamylation levels were quantified using reverse-phase liquid chromatography mass spectrometry (LC-MS; Methods, Extended Data Fig. 1). TTLL7 is a neuronally expressed tubulin glutamylase that preferentially modifies β-tubulin^51,52^. Its expression closely tracks the increase in β-tubulin glutamylation that accompanies neuronal maturation^51^. Total internal reflection (TIRF) microscopy with unmodified microtubules revealed that, at low tau concentrations (< 5 nM), tau partitions between two populations: mobile molecules in low-density regions, and densely packed condensate islands, consistent with previous observations on brain tubulin^15^. The tau islands expand with increasing tau concentration (Extended Data Fig. 2a) and eventually cover the entire microtubule (Extended Data Fig. 2a). As islands expand to cover the entire microtubule, their intensity increases modestly with glutamylation (Extended Data Fig. 2b-d).

To probe how glutamylation affects KIF1A motility through tau condensates, we imaged fluorescently labeled KIF1A on microtubules fully covered with tau at varying glutamylation levels. For all motor assays, tau was used at 20 nM, a concentration that achieves full, uniform condensate coverage on the microtubule lattice (Extended Data Fig. 2a). On unmodified microtubules, KIF1A rarely bound or moved through tau condensates (Fig. 1b), consistent with previous *in vitro* work with brain-derived tubulin^11^. Notably, brain tubulin preparations likely underestimate neuronal glutamylation levels since more than half of brain tissue mass is contributed by glia^53^-cells with low glutamylation^54^-which dilutes the glutamylation signal relative to that of neuronal processes. Strikingly, glutamylation rescues KIF1A motility: unlike on unmodified microtubules, KIF1A lands and moves processively on glutamylated microtubules covered with tau condensates (Fig. 1b, Supplementary Video 1). Quantification reveals that KIF1A landing rate grows monotonically with glutamylation level, rising from 7 ± 5 μm^-1^min^-1^nM^-1^ (mean ± S.D.) on unmodified microtubules to 22 ± 9 μm^-1^min^-1^nM^-1^ at an average of 3 glutamates per β-tubulin (<n_E_>_β_ ∼ 3), and reaching as much as 37 ± 13 μm^-1^min^-1^nM^-1^ on microtubules with <n_E_>_β_ ∼ 6 – a roughly five-fold stimulation – after which the landing rate plateaus (Fig. 1c). Mass spectrometry shows that the preponderance of glutamylated β-tubulin from brain tissue carries a mean of ∼ 6 glutamates^55,56^, the point at which KIF1A landing rates plateaus. Ciliary microtubules – the most glutamylated structures known – carry as many as 20 and 17 glutamates on α-tubulin and β-tubulin tails, respectively^57,58^. These values likely underestimate glutamylation levels in cilia as highly electronegative species are challenging to detect by mass spectrometry. Unlike the landing rate, KIF1A run length does not plateau but continues to increase steadily with glutamylation, reaching 0.5 μm at <n_E_>_β_ ∼ 6 and nearly 1 μm at <n_E_>_β_ ∼26, (Fig. 1d), a ∼ 3-fold increase over the run length at <n_E_>_β_ ∼ 3 (0.4 *vs.* 0.9 μm; Fig 1c), indicating that these two motility parameters respond differently to the glutamylation state of the microtubule.

Detailed analysis of KIF1A motility (Method details) revealed that motor tracks comprise a mixture of processive runs, biased diffusion, and stationary binding events (Figs. 1b, e). Notably, the proportion of stationary binding events decreases steadily with glutamylation, accompanied by an increase in processive and biased runs (Fig. 1e). This progressive shift toward directed motility is captured by the increasing curvature of mean square displacement (MSD) curves with glutamylation level (Fig. 1f) and quantified as the effective velocity (Method details; Fig. 1g). While the speed of individual processive runs does not vary markedly with glutamylation (Fig. 1h), effective velocity increases progressively with glutamate number - from close to zero on unmodified microtubules to ∼ 0.6 μm/s at <n_E_>_β_ ∼ 6, and 0.9 μm/s at <n_E_>_β_ ∼ 26 (Fig. 1g), indicating a greater contribution from longer processive runs to overall motor displacement. In contrast, the rare KIF1A molecules that land on unmodified microtubules are primarily stationary, and the few that do move exhibit only short runs (Figs. 1b, c and d). Notably, the glutamylation-dependent increase in KIF1A motility occurs despite a concurrent modest increase in tau condensate density on the microtubule lattice (Fig. 1i), indicating that the observed enhancement in KIF1A motility is not due to a simple reduction in the tau barrier.

The stimulatory effect on KIF1A landing rates is also seen with tubulin primarily glutamylated on α-tubulin by TTLL6^29,59^. KIF1A landing rate in tau condensates increases monotonically from 6 ± 6 μm^-1^min^-1^nM^-1^ (mean ± S.D.) to 20 ± 12 for unmodified and microtubules with <n_E_>_α_ ∼ 6.1, respectively, and reaches as high as 33 ± 11 μm^-1^min^-1^nM^-1^ for microtubule with <n_E_>_α_ ∼ 16.9 (Extended Data Fig. 3a, b). The enhancement in landing rate as a function of glutamate number is similar to that observed for β-tubulin glutamylation (Extended Data Fig. 3b and Fig. 1c). However, unlike for β-tubulin glutamylation, most landing events result in diffusive motion (Fig. 1f, g and Extended Data Fig. 3c) as reflected by the effective velocity which is close to zero at <n_E_>_α_ ∼ 6.1 and only 0.1 μm/s at <n_E_>_α_ ∼ 16.9, more than 6-fold lower than that on microtubules with <n_E_>_β_ ∼ 13.1 (Fig. 1g). Thus, while glutamylation on either α-or β-tubulin allows KIF1A access to the microtubule through tau condensates, β-tubulin glutamylation is specifically required for processive motility.

Given that KIF1A and KIF5B share overlapping binding sites with tau on the microtubule (Fig. 1a) and yet belong to distinct kinesin families with different mechanochemical properties, we asked whether glutamylation similarly rescues KIF5B motility through tau condensates. We find that in contrast to KIF1A, glutamylation has only a small stimulatory effect on the landing rate of KIF5B on microtubules covered with tau condensates (Fig. 2a, b). Furthermore, the rare KIF5B molecules that land on the microtubule are only stationary or diffusive, and cannot move processively through tau condensates, regardless of glutamylation state (Fig. 2a). Therefore, the two major anterograde motors KIF1A and KIF5B respond to glutamylation in a fundamentally different manner, revealing that the glutamylation code is read in a motor-specific fashion.

**Fig. 2.**
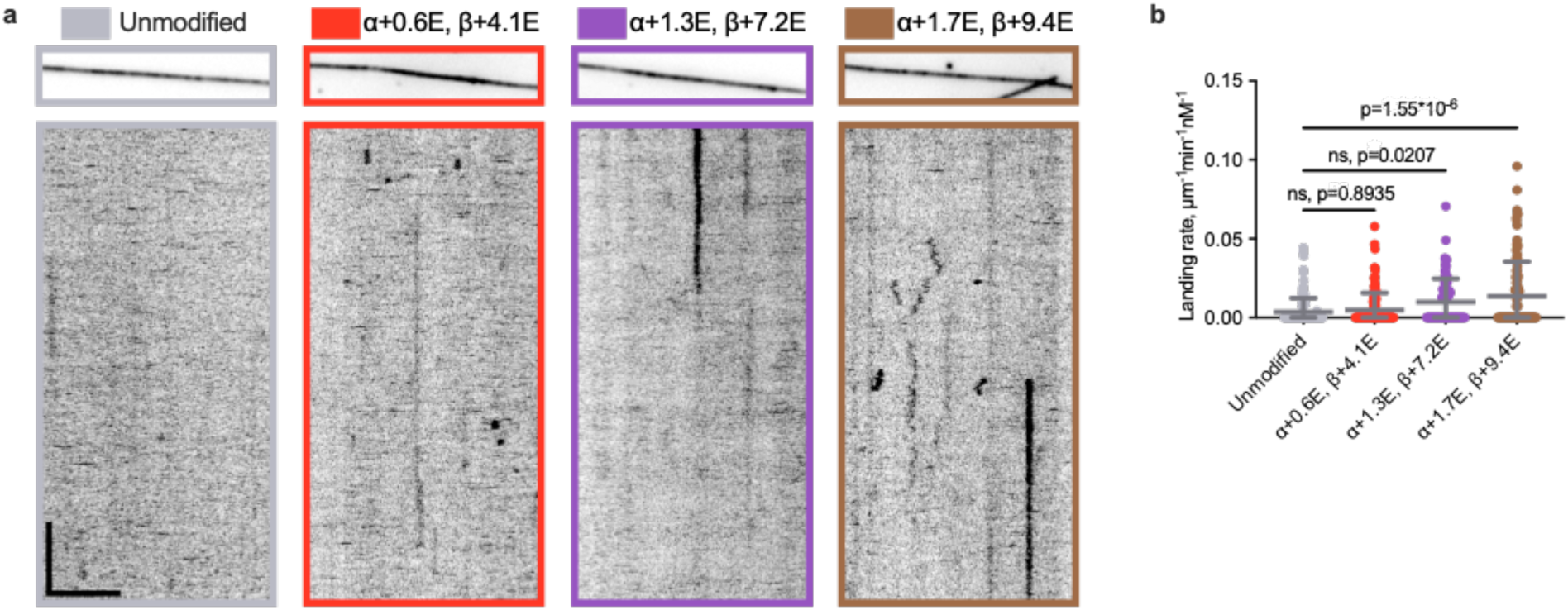
Tau condensates are impenetrable to Kinesin-1 KIF5B regardless of tubulin glutamylation. **a**, Kymographs of KIF5B-GFP motility through tau condensates on unmodified microtubules and microtubules with increasing β-tubulin glutamylation levels. 20 nM sfGFP-tau. Scale bars: horizontal, 5 μm; vertical, 5 s. **b**, KIF5B landing rate, n = 122, 122, 68, and 101 microtubules for unmodified, <n_E_>_β_ ∼ 4.1, 7.2 and 9.4, respectively; p-values determined by one-way ANOVA. 3 independent experiments for all conditions. Data points colored according to glutamylation levels, defined in (**a**).

### Motor-specific K-loop gates motility through tau condensates

Kinesin-3 family members possess a uniquely long and highly positively charged loop in their motor domains, termed the K-loop^60^, that acts as a diffusible tether anchored to the tubulin C-terminal tails to supports high processivity^60–63^. When mutated in KIF1A, both the landing rate and processivity of the motor drop dramatically^64,65^. The KIF1A K-loop has 6 lysine residues, yielding an isoelectric point of 10 - the highest among mammalian kinesins. In contrast, KIF5B has a short K-loop (Fig. 3a). The K-loop of KIF1A is adjacent to the β-tubulin C-terminal tail^63^, making it a compelling candidate for mediating a specific interaction with glutamylated β-tubulin that drives KIF1A motility through tau condensates. To test this hypothesis, we generated a chimeric KIF1A mutant in which the K-loop is replaced with that from KIF5B and examined its motility in the presence and absence of tau and glutamylation. As previously reported^66^, this K-loop substitution reduces the landing rate of the KIF1A chimera mutant drastically (∼ 50-fold; Extended Data Fig. 4a, b). We therefore compensated by performing experiments at higher mutant motor concentrations (13 nM *vs.* 0.1 nM for mutant *vs.* wild-type) to enable comparison with wild-type behavior (Extended Data Fig. 4b). Even at these high mutant motor concentrations, we did not observe landing events on microtubules covered with tau condensates, even if the microtubules are glutamylated (Fig. 3b, c). In contrast, at the same glutamylation level, wild-type KIF1A moves through a mixture of diffusive and processive runs (Fig. 3b) with a mean run length of 0.6 μm (Extended Data Fig. 4c). Together, these data establish the motor-class specific K-loop is necessary both for KIF1A microtubule binding and processive motility through tau condensates on glutamylated microtubules. Interestingly, even in the absence of tau, K-loop loss is more detrimental on glutamylated microtubules than on unmodified microtubules (Extended Data Fig. 4a, d, and e), suggesting that the electronegative glutamylated tails present a steric hindrance to the motor that the KIF1A-specific K-loop uniquely mitigates, and that this interaction is a prerequisite for the full stimulatory effect of glutamylation on KIF1A motility through tau condensates.

**Fig. 3.**
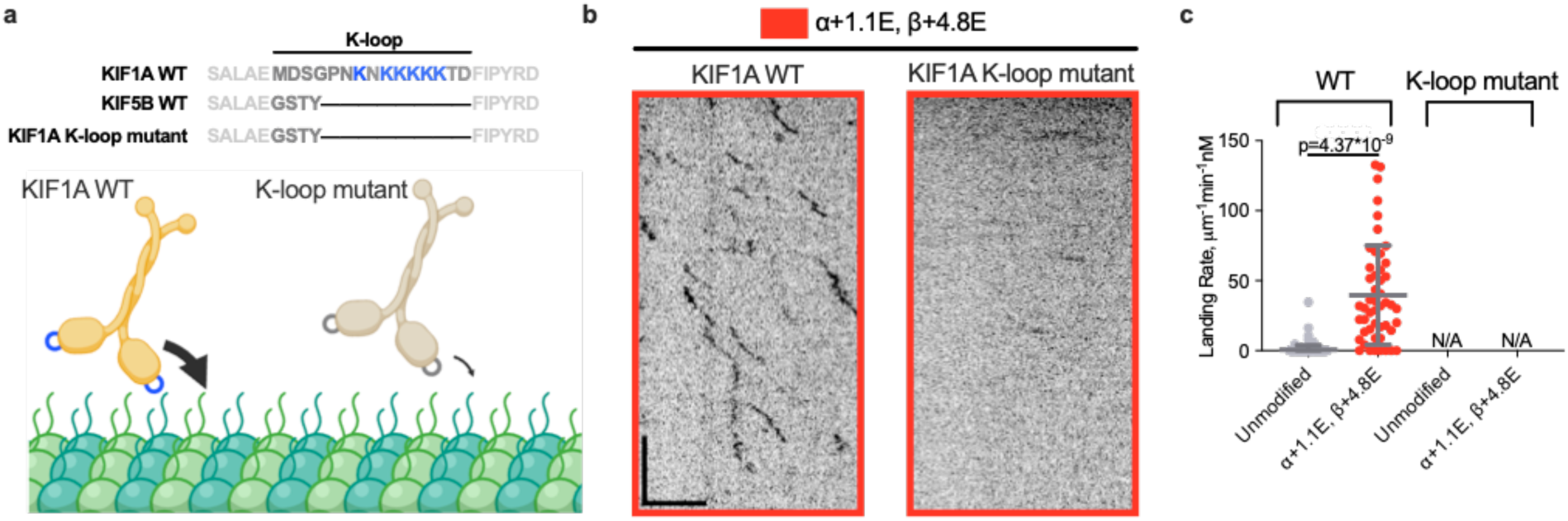
Motor specific K-loop enables kinesin-3 KIF1A motility through tau condensates. **a**, K-loop sequences of KIF1A, KIF5B and the K-loop mutant KIF1A construct. **b**, Kymographs of KIF1A K-loop mutant motility on glutamylated microtubules covered in tau condensates. 20 nM sfGFP-tau. **c**, Landing rate of KIF1A wild-type (WT) and the K-loop mutant on unmodified and glutamylated microtubules covered in tau condensates. 20 nM sfGFP-tau; p-value determined by Wilcoxon rank sum test; WT, n = 133 and 49 microtubules for unmodified and <n_E_>_β_ ∼ 4.8, K-loop mutant, n = 87 and 50 microtubules for unmodified and <n_E_>_β_ ∼ 4.8 conditions. 3 independent experiments for all conditions. Average number of glutamates added to α-and β-tubulin indicated as α + <n_E_>, β + <n_E_>. Data on unmodified and glutamylated microtubules colored grey and red, respectively.

### Glutamylation acts on kinesin motility through tau

Since the KIF1A K-loop sits at the motor-microtubule interface and contacts both tubulin tails^63^, it is well positioned to sense the electronegative glutamates added posttranslationally. We therefore tested whether glutamylation changes the intrinsic motility parameters of KIF1A and KIF5B – independent of tau – as a potential mechanism underlying their differential transit through tau condensates. Surprisingly, KIF1A run length and landing rate change minimally with glutamylation (Fig. 4a, b, and c, Supplementary Video 2). We observed only a small downward drift in speed with β-tubulin glutamylation (Fig. 4d). Similarly, α-tubulin glutamylation has no effect on KIF1A run length and landing rate (Extended Data Fig. 5a, b, and c) and leads only to a small decrease in speed at high glutamylation levels (Extended Data Fig. 5d). The minimal effects of glutamylation on motility parameters are consistent with most previous studies^13,67–69^, though at variance with one report that attributed modest effects of glutamylation to KIF1A run length and pause frequency^69^.

**Fig. 4.**
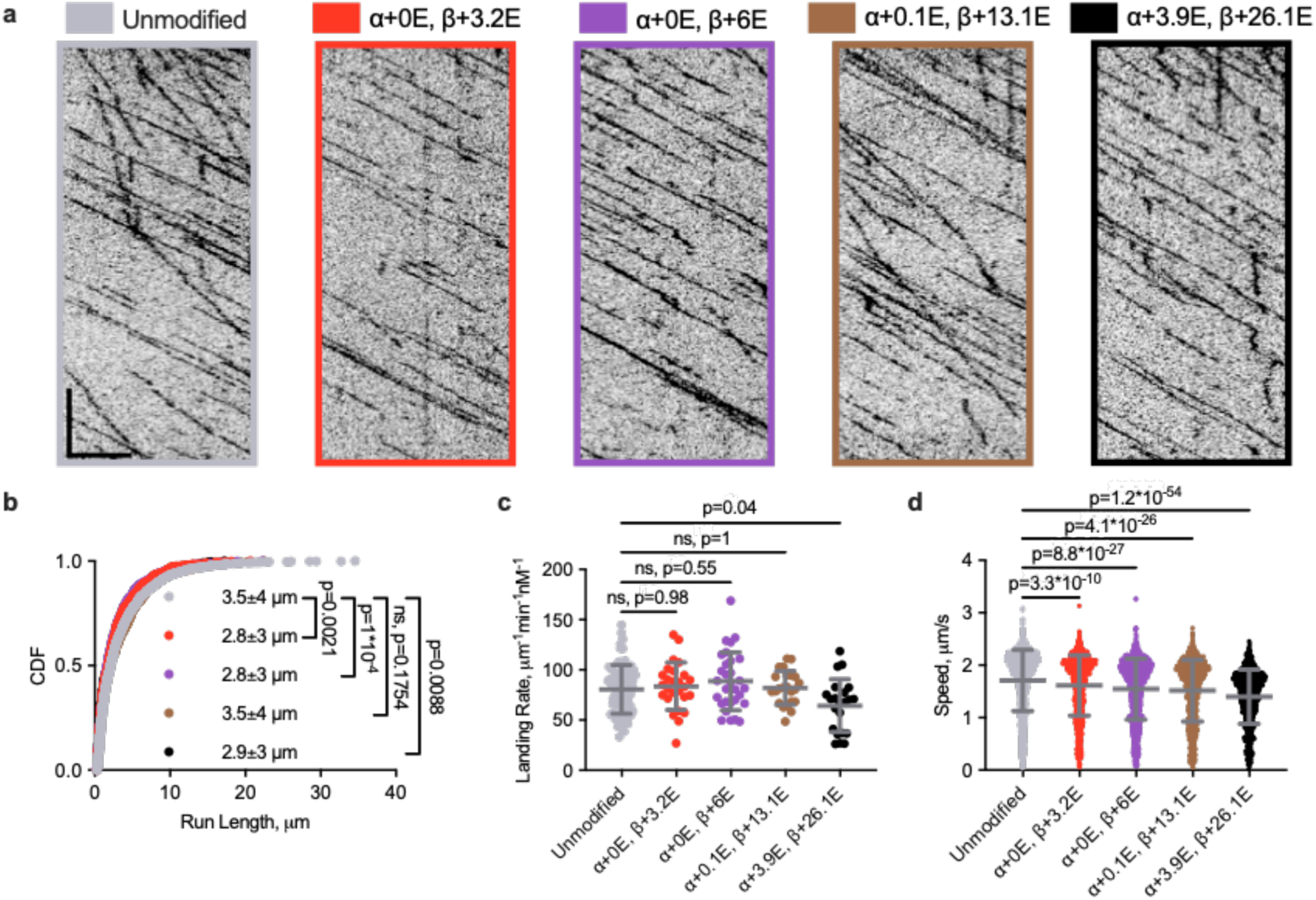
Glutamylation on β-tubulin has no direct effect on kinesin-3 KIF1A motility on tau-free microtubules. **a**, Kymographs of KIF1A-mScarlet motility at increasing β-tubulin glutamylation levels. Scale bars: horizontal, 5 μm; vertical, 5 s. **b**, CDF of KIF1A run length on differentially glutamylated microtubules, mean values ± S.D. shown next to the graph; n = 2263, 698, 929, and 486 molecules for unmodified, <n_E_>_β_ ∼ 3.2, 6, 13.1 and 26.1, respectively; p-values determined by Kolmogorov-Smirnov test. **c**, KIF1A landing rate, ns – not significant, p-values determined by one-way ANOVA, n = 78, 26, 30, 23, and 22 microtubules for unmodified, <n_E_>_β_ ∼ 3.2, 6, 13.1, and 26.1, respectively. **d**, KIF1A speed, n = 3617, 1104, 1227, and 867 molecules for unmodified, <n_E_>_β_ ∼ 3.2, 6, 13.1, and 26.1, respectively; mean values ± S.D. for KIF1A speed were obtained from single gaussian fits, ns – not significant, p-values determined by a two-sided Wilcoxon rank sum test. 12 independent experiments for unmodified and 3 for <n_E_>_β_ ∼ 3.2, 6, 13.1 and 26.1 microtubule conditions. Average number of glutamates added to α-and β-tubulin, α + <n_E_>, β + <n_E_> throughout. Data points colored according to glutamylation levels, defined in (**a**). For microtubule mass spectra see Extended Data Fig. 1. For motility, see Supplementary Video 2.

**Fig. 5.**
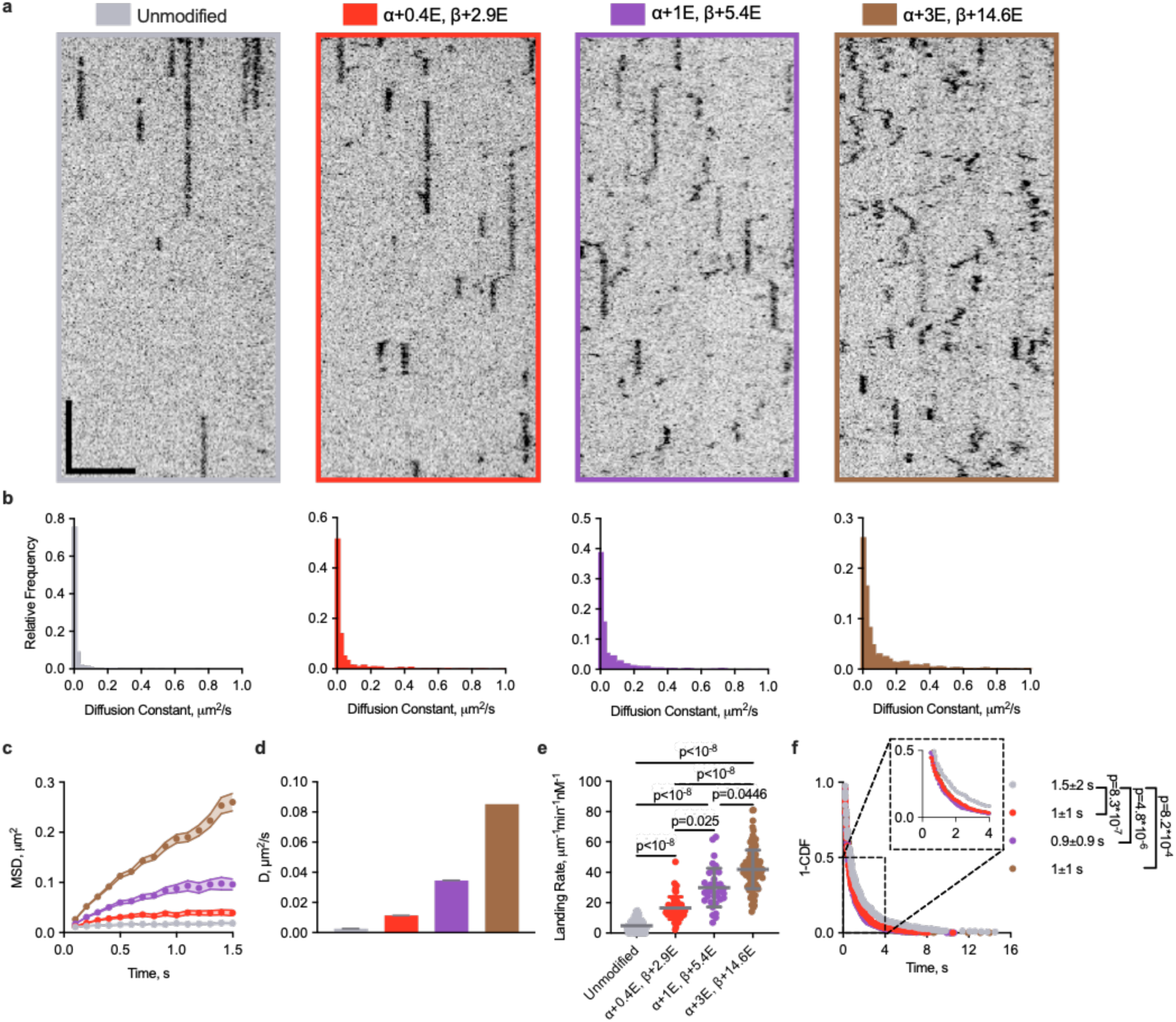
Tubulin glutamylation increases tau condensate diffusivity. **a**, Kymographs of single mScarlet-tau diffusion within tau condensates on microtubules with increasing glutamylation levels. Scale bars: horizontal, 5 μm; vertical, 5 s. 200 pM mScarlet-tau and 50 nM sfGFP-tau **b**, Distributions of tau diffusion constants as a function of tubulin glutamylation. **c**, MSD of single tau molecules in tau condensates as a function of time, n = 1239, 1059, 1353, and 3000 molecules for unmodified, <n_E_>_β_ ∼ 2.9, 5.4, and 14.6, respectively. Shaded areas show S.D. For details see Methods. **d**, tau diffusion constant extracted from MSD fits in (**c**). **e**, Landing rate of single tau molecules within tau condensates, p-values determined by one-way ANOVA, n = 137, 55, 37, and 122 microtubules for unmodified, <n_E_>_β_ ∼ 2.9, 5.4, and 14.6, respectively. **f**, Inverted CDF of dwell times of single tau molecule in tau condensates; mean values ± S.D. shown next to the graph; n = 821, 880, 1182, 2652 molecules for unmodified, <n_E_>_β_ ∼ 2.9, 5.4, and 14.6, respectively; p-values determined by Kolmogorov-Smirnov test. The inset shows a close-up view of the CDFs. 3 independent experiments for all conditions. Average number of glutamates added to α-and β-tubulin, α + <n_E_>, β + <n_E_> throughout. Data points colored according to glutamylation levels, defined in (**a**). See Supplementary Video 4.

Glutamylation likewise has no direct effect on KIF5B run length or landing rate (Extended Data Fig. 6a, b and c, Supplementary Video 3). The only effect is a slight reduction in speed with glutamylation (Fig. 6d). This reduction in speed for both KIF1A and KIF5B is consistent with an increased steric hindrance due to the glutamate chains on the tails. *In toto*, our experiments establish that KIF1A and KIF5B are largely insensitive to glutamylation on MAP-free microtubules and that the differential motility of these motors through tau condensates does not arise from intrinsic glutamylation sensitivity of the motor.

**Fig. 6.**
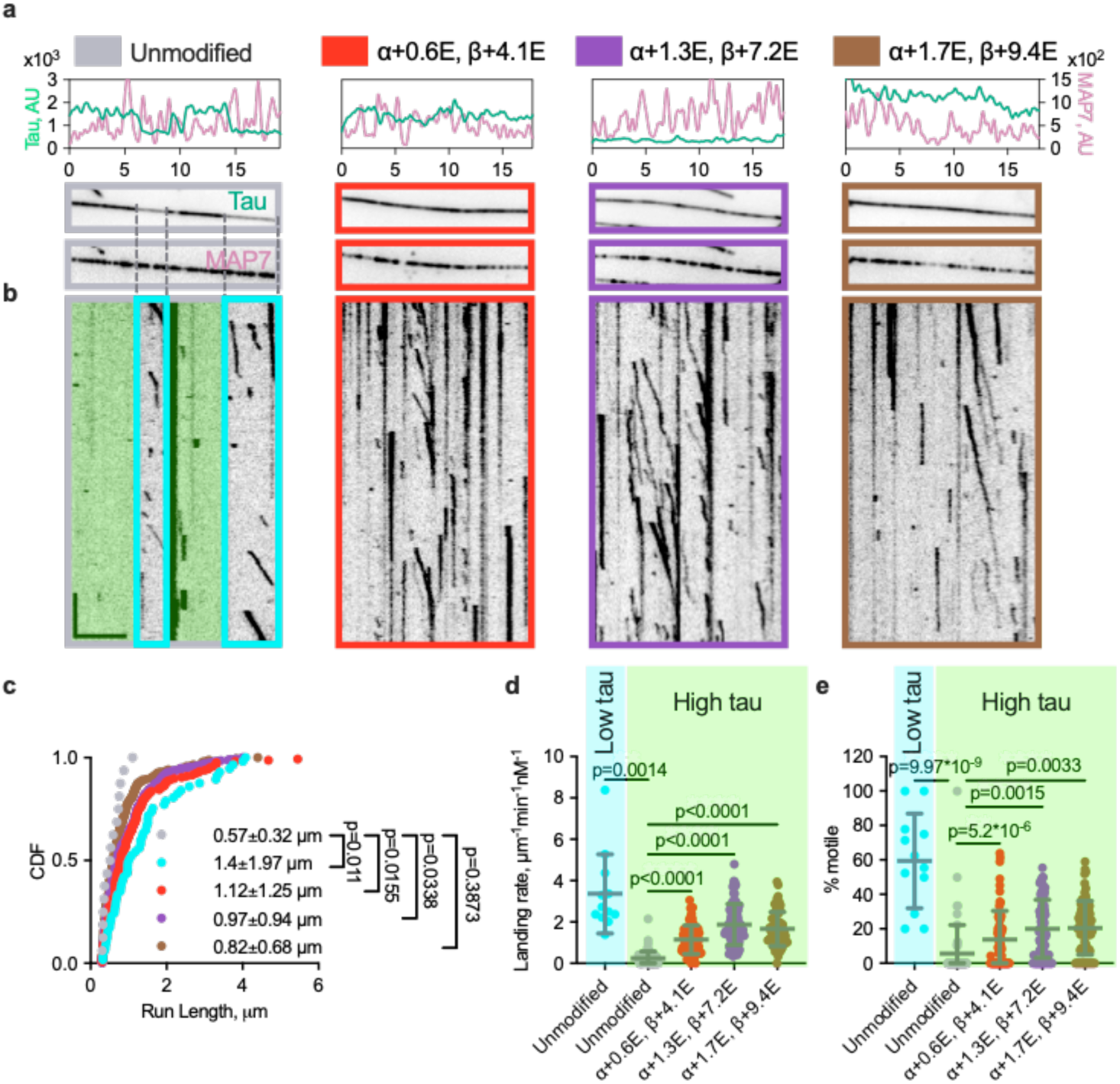
Glutamylation promotes MAP7-tau mixing but does not support high processive motility even in the presence of MAP7. **a**, Glutamylation promotes tau-MAP7 mixing. Plots show a line scans of tau (green, left y-axis) and MAP7 intensity (pink, right y-axis) on the microtubule shown below. 1 nM LD655-labeled MAP7, 20 nM sfGFP-tau. **b**, Kymographs of KIF5B-GFP motility on differentially glutamylated microtubules shown above in panel (**a**). Scale bars: horizontal, 5 μm; vertical, 5 s. **c**, CDF of run lengths of motile KIF5B at increasing glutamylation levels, mean values ± S.D. shown next to the graph; n = 57, 16, 167, 350, 258 molecules in low tau and high tau density regions (condensates) on unmodified microtubules, and in tau condensates (high density tau regions) on glutamylated microtubules with <n_E_>_β_ ∼ 4.1, 7.2, 9.4, respectively. **d**, KIF5B landing rate, n = 12 and 72 regions in low tau and high tau density regions (condensates) on unmodified microtubules, respectively, and n = 70, 81, and 68 microtubules for <n_E_>_β_ ∼ 4.1, 7.2, 9.4, respectively. **e**, Percentage of motile KIF5B motors at increasing glutamylation levels; n = 12 and 54 regions in low tau and high tau density regions (condensates) on unmodified microtubules, respectively, and n = 70, 81, and 68 for tau condensates on microtubules with <n_E_>_β_ ∼ 4.1, 7.2, and 9.4, respectively. p-values determined by one-way ANOVA test in (**d**), (**e**), and by Kolmogorov-Smirnov test in (**c**). 3 independent experiments for all conditions. Average number of glutamates added to α-and β-tubulin, α + <n_E_>, β + <n_E_> throughout. Data points colored according to glutamylation levels, defined in (A). See Supplementary Video 5.

### Glutamylation increases tau diffusivity within condensates

Given that both KIF1A and KIF5B are insensitive to glutamylation in the absence of tau, we turned our attention to the effects of tubulin glutamylation on tau. Tau condensates are dense, ∼ 0.8 tau molecules per tubulin dimer^16^. Therefore, the motor is severely restricted by the lack of microtubule binding sites it can access within the condensate. Since tau interacts with the tubulin tails which together with the posttranslationally added glutamate chains form a dense electronegative polymer brush, we investigated whether tubulin glutamylation alters the dynamics of tau molecules within condensates. To directly probe how glutamylation affects tau condensate dynamics, we tracked single mScarlet-tau molecules within GFP-tau condensates on microtubules at varying glutamylation levels (Methods). On unmodified microtubules, tau molecules are predominantly stationary (Fig. 5a). Glutamylation progressively increases the mobility of tau within the condensate: the mobile tau fraction increases from 17% on unmodified microtubules to 37%, 47% and 59% for <n_E_>_β_ of 2.9, 5.4 and 14.6, respectively (Fig. 5a, b, Supplementary Video 4). Consistent with this, the diffusion constant of tau within condensates increases steadily with tubulin glutamylation, reaching a 43-fold enhancement at <n_E_>_β_ ∼14.6 compared to unmodified microtubules (Fig. 5c, d). Notably, glutamylation has the opposite effect on diffusive tau – the low-density, non-condensate population observed at lower concentrations (5 nM or below; Extended Data Fig. 2) – whose diffusivity on the microtubule lattice decreases modestly with glutamylation (Extended Data Fig. 7a, b). The different response to glutamylation of condensate-associated *versus* sparsely bound tau suggests different modalities of microtubule engagement, multivalent interactions with condensates *versus* individual binding at sparse densities. The increase in tau diffusion within condensates is accompanied by a dramatic, up to 8-fold increase in landing rate on microtubules with <n_E_>_β_ ∼14.6 *versus* unmodified ones (Fig. 5e), as well as reduction in dwell times on the microtubule (Fig. 5f). Together, the increased tau diffusivity and faster exchange between the microtubule-bound and solution states effectively fluidize the tau condensates. Notably, tau dwell time changes only modestly between <n_E_>_β_ ∼ 2.9 and 14.6 (1.5 s versus 1 s, Fig. 5f), while tau diffusivity increases ∼ 30-fold over the same range (Fig. 5d), closely tracking the continued increase in KIF1A landing rate and processivity (Fig. 1c, d. This correlation identifies tau diffusion within the condensates, rather than tau turnover, as the dominant driver of glutamylation-dependent KIF1A motility.

### Glutamylation promotes MAP7-tau mixing but is not sufficient for KIF5B motility

MAP7 acts as a KIF5B-specific co-factor that bridges the motor and the microtubule, increasing both the landing rate and run length of KIF5B^11,70–72^. We therefore investigated whether the MAP7 co-factor can help KIF5B overcome the inhibitory effect of tau on motility. MAP7 and tau were previously reported to demix on microtubules^17^, setting up exclusionary zones that have one MAP and not the other. Consistent with this, on unmodified microtubules, we see an anti-correlation between MAP7 and tau intensity, indicating that both proteins favor self-association under these conditions (Fig. 6a). Consequently, KIF5B motility is supported in MAP7 dense regions, but not in tau-dense regions (Fig. 6a, b, Supplementary Video 5). Glutamylation promotes mixing of these two MAPs (Fig. 6a), consistent with the increased tau mobility described above (Fig. 5). This mixing increases KIF5B landing rate to microtubule regions with high tau density (Figs. 6b-d, Supplementary Video 5), which in the absence of MAP7 is almost ∼100-fold lower (Extended Data Fig. 6b). However, despite the enhanced microtubule access, only a small fraction of motors initiate processive runs: 5, 14, and 20% for <n_E_>_β_ ∼ 4.1, 7.2, and 9.4, respectively (Fig. 6e). On tau-free microtubules MAP7 supports long KIF5B processive runs exceeding 10 μm, (Extended Data Fig. 8a, Supplementary Video 6), and increases the motor landing rate four orders of magnitude from ∼ 0.1 to 1000 μm^-1^*min^-1^*nM^-1^ (Extended Data Figs. 6b and 8b), independently of tubulin glutamylation. Glutamylation has no additional stimulatory effect on the KIF5B-MAP7 complex; its only influence is a slight reduction in speed on glutamylated microtubules (Extended Data Fig. 8c), as observed for KIF5B alone (Extended Data Fig. 6d). In contrast to tau, MAP7 dynamics on the microtubule are insensitive to glutamylation (Extended Data Fig. 9a, b, and c, Supplementary Video 7). Thus, although MAP7 increases KIF5B access through tau condensates, it fails to support robust processive motion even in the presence of glutamylation, underscoring the motor-and MAP-specific nature of glutamylation-dependent transport regulation. In contrast, the KIF5B adapter TRAK1 (the Trafficking Kinesin Protein 1) which links kinesin to mitochondria, supports KIF5B motility through tau condensates once motility is initiated^73^, but unlike MAP7 is unable to facilitate KIF5B access to these islands.

### MAP9 - glutamylation synergy enables high KIF1A processivity

KIF1A also has a motor-specific co-factor, MAP9, which increases its microtubule landing rate^11,74^. Given the failure of MAP7 to rescue KIF5B motility through tau condensates, we asked whether MAP9 could further potentiate KIF1A motility through tau condensates in the presence of glutamylation. We find that on unmodified microtubules covered with tau, KIF1A binds only rarely and is primarily non-motile, regardless of MAP9 concentration (Figs. 7a, b, and c, Extended Data Fig. 10a-i), indicating that the MAP9 tether cannot overcome the cohesive barrier imposed by tau condensates in the absence of tubulin glutamylation. In striking contrast, on glutamylated microtubules MAP9 dramatically increases both KIF1A landing rate and run length through tau condensates. This stimulation is MAP9 concentration dependent (Fig. 7b and c, Extended Data Fig. 10b-i, Supplementary Videos 8 and 9). Both landing rate and run length increase ∼ 2-fold at 0.5 nM MAP9 and ∼ 3-fold at 1 nM MAP9, relative to the MAP9-free conditions (Fig. 7b and c). Strikingly, at 1 nM MAP9, the KIF1A landing rate on glutamylated tau-covered microtubules exceeds that of the motor alone on tau-free microtubules (Figs. 7b and 4c), therefore reversing the inhibitory effects of tau. Moreover, MAP9 stimulation within tau condensates at 1 nM MAP9 is significantly higher (3-fold) than on tau-free microtubules where even saturating MAP9 concentrations (6-60 nM) achieve a maximal increase in landing rate of less than 50%^74^. *In toto* these data demonstrate that MAP9 acts synergistically with glutamylation to drive high-volume KIF1A traffic on microtubules covered with tau, in contrast to KIF5B where MAP7 and glutamylation cannot overcome the inhibitory effects of tau (Figs. 7d and e).

**Fig. 7.**
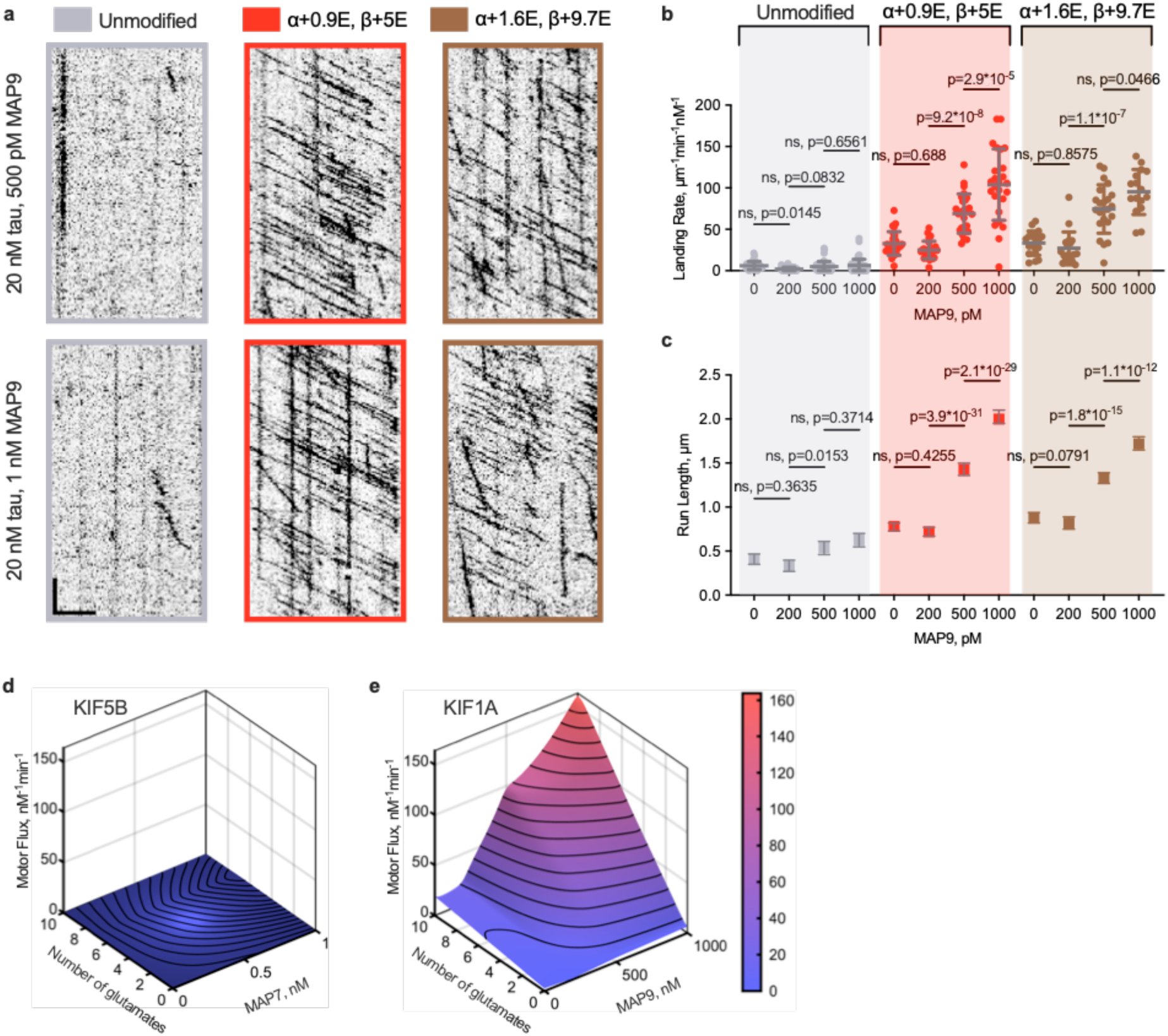
Glutamylation synergizes with MAP9 to facilitate robust KIF1A motility on microtubules covered in tau condensates. **a**, Kymographs of KIF1A-mScarlet motility on differentially glutamylated microtubules covered in tau condensates in the presence of 0.5 nM (top) and 1 nM MAP9; 20 nM sfGFP-tau concentration. **b**, KIF1A-mScarlet landing rate on differentially glutamylated microtubules covered in tau condensates at different MAP9 concentrations, n = 54, 25, and 16 microtubules for unmodified, <n_E_>_β_ ∼ 5 and 9.7, respectively. **c**, Mean KIF1A run lengths on differentially glutamylated microtubules covered in tau condensates at different MAP9 concentrations, n = 307, 2639, and 2367 molecules for unmodified, <n_E_>_β_ = 5 and 9.7, respectively; error bars show 95% confidence intervals. **d-e**, Motor flux of KIF5B **(d)** and KIF1A **(e)** through tau condensates as a function of glutamylation and concentration of their respective activators, MAP7 **(d)** or MAP9 **(e)**. Color bar on the right in **(e)** shows motor flux level. Motor flux determined as detailed in Methods. Scale bars in (**a**): horizontal, 5 μm; vertical, 5 s; p-values determined by one-way ANOVA test in (**b**) and by Kolmogorov-Smirnov test in (**c**). 6 independent experiments for unmodified and 3 for <n_E_>_β_ ∼ 5, and 9.7 microtubule conditions. Average number of glutamates added to α-and β-tubulin, α + <n_E_>, β + <n_E_> throughout. Data points colored according to glutamylation levels, defined in (**a**). See Supplementary Videos 8 and 9.

## DISCUSSION

Our work establishes several broad principles to advance our understanding of transport in the crowded MAP-dominated neuronal microtubule environment as well as the biophysics of microtubule-associated MAP condensates. First, tubulin glutamylation, highly abundant in neuronal processes, fluidizes condensates formed by tau by increasing tau diffusivity dramatically (more than 30-fold) and accelerating tau exchange between the microtubule-bound and solution states, therefore remodeling tau condensates from impassible barriers to structures permissive to motor transit. This remodeling is mechanistically distinct from a simple reduction in tau occupancy to allow motor transport, as tau density on the microtubule increases with glutamylation. We propose that glutamylation fluidizes or increases disorder in the condensate by disrupting polyvalent homotypic tau interactions through steric and electrostatic repulsion imposed by the disordered, electronegative polyglutamate chains on the tubulin tails. The mechanism we uncover, that tubulin glutamylation changes the dynamics of a MAP condensate to regulate motor access, is likely not limited to tau. Other neuronal MAPs that form microtubule-associated condensates^75^ share common mechanisms of microtubule recognition, suggesting that glutamylation-induced fluidization may be a more general strategy to tune the motile landscape across diverse MAP-decorated microtubule domains.

Second, we show that selective motor transport is not hardwired but emerges from the synergy of motor properties, MAP identity and the tubulin code. Specifically, we show that tau condensate properties synergize with intrinsic motor properties to support selective motility of KIF1A, but not that of KIF5B (Figs. 7d, e and Extended Data Fig. 11). Kinesin motors alternate binding of their two motor domains to achieve processive motion on the microtubule. Therefore, the binding and unbinding kinetics of the motor are key to its ability to navigate the dynamic obstacles course formed by tau on glutamylated microtubules. KIF1A adopts a two-head-bound state, which minimizes detachment, leading to predominantly 16 nm steps with occasionally longer steps^76^. Unlike KIF5B, KIF1A can stay attached to the microtubule in the ADP-bound state for tens of seconds^76–79^, longer than the average dwell time of tau molecules within condensates on glutamylated microtubules (∼ 1 s, Fig. 5f). The extended attachment time increases the probability that tau diffuses to its next binding site or leaves the microtubule altogether, freeing a binding site for the motor. An additional advantage of KIF1A over KIF5B may lie in its ability to sidestep between protofilaments^80^, providing alternative binding site options to navigate around tau-occupied regions, whereas KIF5B is confined to a single protofilament track^81,82^. Furthermore, KIF1A can quickly rebind once it lets go^83–85^. We hypothesize that together these intrinsic KIF1A-specific properties ensure that the motor has a lower probability of diffusing away from the microtubule while waiting for a tau molecule to move out of the way so that it can take the next step or rebind. Consistent with such a model, Kip3, a super-processive yeast kinesin which can pause without detaching when a binding site is unavailable^86^ can traverse tau condensates^16^. Interestingly, while glutamylation on either α-or β-tubulin enhances kinesin access to tau condensates, processive motion is supported only by β-tubulin glutamylation. This implies that tau enables regioselective interaction of KIF1A with glutamylated tubulin tails to support processive motility. Adapters such as MAP7 and MAP9 provide additional tethering points to the microtubule for KIF5B and KIF1A, respectively; however, their effects within tau condensates are very different. Even in the presence of MAP7, KIF5B runs are still severely hindered in tau condensates indicating that intrinsic motor properties are determinant. In contrast, MAP9 dramatically potentiates KIF1A flux through tau condensates (Figs. 7d and e). Strikingly, this stimulation is stronger than the maximal stimulation achieved for KIF1A with MAP9 on tau-free microtubules^74^, an unexpected finding suggesting that MAP9 functions optimally to potentiate KIF1A motility in the crowded tau condensate environment. Critically, however, MAP9 cannot support KIF1A motility at any concentration tested, establishing that the observed robust processive transport through tau condensates requires the obligate synergy between glutamylation and MAP9 (Fig. 7e, Extended Data Fig. 11c). For KIF5B, Trak1 may be required to help overcome intrinsic motor limitations within tau-rich environments, with robust processive transport through tau condensates requiring the concerted action of glutamylation, MAP7 and Trak1.

Third, our multilayered reconstitution system that integrates MAPs and tubulin modifications, the first in the field, demonstrates the importance of experimentally addressing complex macromolecular interactions because these can lead to qualitatively distinct motor behaviors that arise only at the intersection of multiple regulatory layers. Specifically, we show that kinesin motility on crowded tau-covered microtubules is an emergent property that cannot be predicted from any single regulatory input alone, but arises from the combinatorial, motor-specific interplay between MAPs and tubulin glutamylation.

Fourth, while posttranslational modifications, most prominently phosphorylation, have been shown to alter network interactions within protein condensates to modulate their internal dynamics^87^, in many cases increasing their dynamicity^88–90^, our work reveals that glutamylation operates through a fundamentally different mechanism: rather than directly modifying the condensate-forming protein, it acts *in trans* through the microtubule scaffold that nucleates condensation, establishing a new paradigm for how the physical properties of a condensate can be regulated by the underlying polymer template. Beyond tubulin, glutamylation targets a broad range of non-tubulin targets^91–93^, many of which contain large intrinsically disordered regions, raising the possibility that glutamylation may function as a more general regulator of protein dynamics across multiple cellular contexts.

Our findings also carry implications for tauopathies. Tau condensate dysregulation is a feature of Alzheimer’s disease and frontotemporal dementia, and disrupted glutamylation homeostasis is also associated with neurodegeneration. Alongside the well-established roles of tau aggregation in driving transport failure, our demonstration that glutamylation acts on tau condensates through a tubulin-based route opens new avenues for understanding how transport failure arises in tauopathies.

## EXTENDED DATA

Extended Data Figures 1-11

## SUPPLEMENTARY INFORMATION

Supplementary Figures 1-11

Supplementary Videos 1-9

## METHODS

### Protein expression and purification

*Homo sapiens* full-length tau (UNIPROT# P10636-8, 2N4R, aa 1-441), truncated *Rattus norvegicus* KIF1A (UNIPROT# F1M4A4, aa 1-393), and *Homo sapiens* MAP9 (UNIPROT# Q49MG5, aa 1-647) were cloned into pET28A vectors using Gibson assembly. Tau was tagged with Strep-Tag II at the N-terminus, followed by either sfGFP or mScarlet; MAP9 was tagged with Strep-Tag II at the N-terminus, followed by sfGFP; KIF1A was tagged with mScarlet at the C-terminus, followed by Strep-Tag II. sfGFP-tau was prepared as previously described^11^. For bacterial expression of mScarlet-tau, sfGFP-tau and KIF1A-mScarlet BL21 Rosetta 2(DE3) pLysS cells were grown at 37°C until ∼O.D. 0.6 and protein expression was induced with 0.1 mM IPTG. Cells were grown overnight at 16°C, harvested, and frozen. Cell pellets were resuspended in lysis buffer (50 mM Tris pH 8, 200 mM KCl, 2 mM MgCl_2_, 1 mM EGTA, 10% (v/v) glycerol) with protease inhibitor cocktail (Roche #5056489001), 1 mM DTT, 1 mM PMSF, and DNAse I. Cells were lysed by microfluidization in an Emulsiflex press and cleared by centrifugation at 40,000 x g for 1 h. The clarified lysate was passed over a column packed with Streptactin XT Superflow resin (Qiagen/IBA). After 2 h incubation, the column was washed with four column volumes of lysis buffer, then bound proteins were eluted with 50 mM D-biotin (CHEM-IMPEX) in lysis buffer. Eluted tau constructs were further purified on a HiTrap Q HP column in (50 mM Tris pH 7.5, 100 mM NaCl, 2 mM MgCl_2_, 1 mM EGTA, 10% (v/v) glycerol) buffer and eluted with a 100-400 mM NaCl gradient using an Akta pure system. Peak fractions were pooled and concentrated using Amicon concentrators followed by passage through a size exclusion column (Superdex 200 10/300 GL, GE Healthcare). Peak fractions were pooled, concentrated, and flash frozen in liquid nitrogen. KIF1A-mScarlet and the KIF1A-K-loop mutant were affinity purified using a Streptactin XT Superflow column followed microtubule binding and release. Specifically, 200 μl of the streptactin column eluate was thawed and incubated at room temperature for 15 min. Taxol was added to the eluate to a final concentration of 20 μM. 50 μl of 2 mg/ml taxol-stabilized microtubules were then added to the eluate, the mixture was incubated at room temperature for 15 minutes to allow the motors to bind to the microtubules and then passed through a 250 μL glycerol cushion (80 mM PIPES, 2 mM MgCl_2_, 1 mM EGTA, 1 mM DTT, 10 μM taxol, and 60% glycerol, pH 6.8) by centrifugation at 50,000 rpm for 10 minutes at room temperature in a TLA-100 rotor (Beckman). Next, the supernatant was discarded and the pellet was resuspended in 50 μl high-salt release buffer (30 mM HEPES, 300 mM KCl, 2 mM MgCl_2_, 1 mM EGTA, 1 mM DTT, 10 μM taxol, 3 mM ATP). The released motors were then separated from the microtubules by centrifugation at 80,000 rpm for 10 minutes at room temperature. The supernatant containing the active motors was aliquoted and flash frozen in liquid nitrogen.

*Homo sapiens* kinesin-1 KIF5B (UNIPROT# P33176, aa 1-560) with a C-terminal GFP (K560-GFP) and full-length *Homo sapiens* MAP7 (UNIPROT# Q14244-1, aa 1-749) both have an N-terminal ZZ affinity tag followed by a TEV protease cleavage sequence. MAP7 has an N-terminal ybbR sequence. The K560 construct has tandem GFP and SNAP-tags connected by a 1xGS linker sequence at its C-terminus. The K560-SNAP-GFP and MAP7 constructs were purified from baculovirus-infected SF9 cells. A cell pellet from a 1.35 L culture of SF9 cells was thawed in 200 mL lysis buffer (25 mM HEPES pH 7.4, 1 M KCl, 10% glycerol, protease inhibitor (Roche #5056489001), 1 mM PMSF, 1 mM DTT, 0.1 mM ATP). Cells were lysed with a Wheaton glass dounce homogenizer. The lysate was centrifuged for 1h at 65,000 rpm in a Ti70 rotor (Beckman). The supernatant was incubated with 3 mL IgG sepharose resin for 1 h. The resin was then washed with 50 mL lysis buffer followed by 50 mL storage buffer (25 mM HEPES pH 7.4, 300 mM KCl, 1 mM EGTA, 10 mM MgCl_2_, 10% glycerol, 1 mM DTT). 0.1 mM ATP was added to the buffer for the K560 purification). The resin was resuspended in 5 mL storage buffer and transferred to a 5 mL Eppendorf tube. For SNAP-tag labeling, the resin-bound protein was incubated with 10 nmol SNAP-ligand conjugated dye for 1 h on ice. For ybbR-tag labeling, the resin-bound protein was incubated with coenzyme-A conjugated dye (Lumidine Technologies) and 1 μM *Bacillus subtilis* phosphopantetheinyl transferase (SFP) for 30 min at room temperature. The labeled proteins were eluted by addition of 30 μL of 2 mg/ml TEV protease for 30 min at room temperature. The eluted proteins were further purified by size exclusion chromatography on a Superdex 200 10/300 GL (GE Healthcare) column. Peak fractions were collected, pooled, concentrated, and flash frozen in liquid nitrogen.

### Preparation of differentially glutamylated microtubules

Human unmodified tubulin was purified from tSA201 cells by the TOG affinity method^94^ using a TOG1 domain affinity column^95^. Taxol-stabilized unmodified microtubules were prepared by polymerizing purified unmodified human tubulin with 1.5% biotin-labeled porcine brain tubulin (T333P, Cytoskeleton). For control microtubules, we further added 1.5% HiLyte 647-labeled porcine brain tubulin (TL670M, Cytoskeleton) to distinguish unmodified and glutamylated microtubules in motility assays (see next section). The polymerization mix was supplemented with 10 μM taxol after 2 h of polymerization and incubated at 37°C for 2 h. Microtubules were glutamylated as described previously^28^. Briefly, microtubules were glutamylated by incubation with *Xenopus tropicalis* tubulin tyrosine ligase-like 7 (TTLL7, for β-tubulin modification) or *Homo sapiens* TTLL6 (for α-tubulin modification) at a 1:10 molar ratio of enzyme to tubulin in 20 mM HEPES pH 7.0, 50 mM NaCl, 10 mM MgCl_2_, 1 mM Glu, 1 mM ATP, 10 mM taxol at room temperature for various incubation times (1 h for <n_E_> ∼6, 2 h for <n_E_> ∼13, 8 h for <n_E_> ∼26). Control reactions were performed with 1 mM aspartate which is not a substrate for TTLL7 or TTLL6. Enzymes were removed by addition of 0.3 M KCl and sedimentation through a 60% glycerol cushion at 100,000 x g for 12 min at 30°C. Microtubules were stored in BRB80 (80 mM PIPES, 1 mM MgCl_2_, 1 mM EGTA, pH 6.8), 14.3 mM β-mercaptoethanol, 10 μM taxol. The number of glutamates added to α-and β-tubulin was determined by LC-MS^28,95^. The spectra were deconvoluted in MassHunter software (Agilent) and display the characteristic distribution of masses with peaks separated by 129 Da corresponding to one glutamate. The extent of tubulin glutamylation on α-and β-tubulin was quantified by calculating the weighted average of peak intensities for each tubulin species present.

### Tau microtubule binding at different tau concentrations

Solutions of sfGFP-tau at 1, 5, and 20 nM in assays buffer (20 mM HEPES pH 7.4, 40 mM KCl, 2 mM MgCl_2_ 1 mM EGTA, 10% (v/v) glycerol, 0.5% (w/v) pluronic F127, 2 mM ATP, 1 mM TCEP, 0.2 mg/mL casein, and 10 μM taxol) supplemented with an oxygen scavenger mix to remove free oxygen from solution (7.5 U/μL catalase, 0.15 U/μL glucose oxidase, 20 mM D-glucose) were sequentially perfused in the flow chamber with microtubules immobilized on the surface and incubated for 10 min to allow tau to reach equilibrium on the microtubule lattice. Micrographs of the microtubule-bound tau were collected using a 488 nm channel at 100 ms exposure time. The density of tau on microtubules was quantified by dividing the sfGFP-tau intensity (RawIntDen) by the microtubule length in FIJI^96^.

### KIF1A motility assays

Flow chambers for TIRF assays were made of silanized coverslips and slides. Flow chamber construction and silanization protocol is covered in detail in a previously published methods chapter^28,97^. Microtubules prepared as described in the previous section were immobilized in the flow chamber with 0.02 mg/ml NeutrAvidin (Thermo Scientific) and washed with BRB80 supplemented with 20 μM taxol, 14.3 μM β-mercaptoethanol, and 2 mg/ml casein. Unmodified microtubules were present alongside each glutamylated microtubule sample for an internal control. For kinesin-3 motility assays on tau-free microtubules, KIF1A-mScarlet was diluted in the assays buffer (20 mM HEPES pH 7.4, 40 mM KCl, 2 mM MgCl_2_ 1 mM EGTA, 10% (v/v) glycerol, 0.5% (w/v) pluronic F127, 2 mM ATP, 1 mM TCEP, 0.2 mg/mL casein, and 10 μM taxol) supplemented with an oxygen scavenger mix to remove free oxygen from solution (7.5 U/μL catalase, 0.15 U/μL glucose oxidase, 20 mM D-glucose) and perfused into the chamber with microtubules already immobilized on the surface.

For KIF1A in the presence of tau, either 20 nM sfGFP-tau or a mixture of 20 nM sfGFP-tau and 0.2, 0.5, or 1 nM sfGFP-MAP9 in assay buffer was perfused into the chamber with microtubules already immobilized on the surface and incubated for 10 min to allow the chamber to equilibrate. Next, KIF1A-mScarlet was perfused into the chamber at the concentration of interest together with sfGFP-tau and sfGFP-MAP9. Kinesin motility was recorded at a framerate of 5 or 10 Hz for 30-60 s at 100 ms exposure per frame. For the experiments without tau, the same kinesin concentrations and imaging settings were used as those used in the experiments with tau. Kinesin motility was analyzed using FIESTA (v. 1.6)^98^ or FIJI^96^. To obtain motility parameters (speed, run length, and landing rate), the tracking data from FIESTA was exported and analyzed in MATLAB using custom-written scripts (deposited on GitHub). Data analysis in ImageJ was performed manually using the KymographBuilder plugin to make kymographs in the kinesin channel. Speed values were thresholded using a displacement cutoff of 0.3 μm. Run length distributions were fit to a one phase exponential decay, and the decay constant was derived to show an average run length value. KIF1A tracks were categorized into four groups based on their motility: processive, biased, diffusive and stationary. A motility event was considered processive if no more than two backsteps occurred during the run. A motility event was marked as biased if the kinesin molecule performed multiple (>2) backsteps and its displacement was larger than 0.5 μm. Events were marked as diffusive if they had random motility with a net displacement smaller than 0.5 μm, or stationary if the displacement was less than 0.3 μm. Landing rate was quantified by dividing the number of molecules that interact with a microtubule by a product of the microtubule length, total measurement time, and the soluble tau concentration. Run lengths were extracted from the tracking data. Only run lengths of molecules whose entire path can be observed during imaging were counted, i.e., molecules that landed on a microtubule before the beginning of imaging or dissociated after its end were not taken into account. For quantification of speeds, only tracks with a displacement more 0.3 μm were taken into account to discard occasional stationary molecules. The full-width-at-half-maximum of the point spread function of stationary single dyes in our setup is ∼300 nm, which was taken as a cutoff to classify motile particles for speed measurements. Mean squared displacement analysis was performed using a built-in function in FIESTA. The values were plotted in GraphPad Prism 9 and fit to a quadratic one-dimensional diffusion equation with a directed motion term, MSD = v^2^τ^2^ + 2Dτ + C (samples with high glutamylation levels, <n^E^>_β_ > 6), or to a linear one-dimensional diffusion equation, MSD = 2Dτ + C (samples with low glutamylation levels <n^E^>_β_ ≤ 6), where τ is a lag time, v – is an effective velocity, D is a diffusion constant, and C is an offset. Motor flux is calculated as a product of mean run length and landing rate of a motor within tau condensates. Three-dimensional surfaces showing motor flux as a function of glutamylation and activator concentration (MAP9 or MAP7) were created in MATLAB using normalized gaussian smoothing applied to a natural neighbor interpolation grid.

### KIF5B motility assays

Flow chambers were prepared as described in the previous section. For kinesin-1 motility assays in the presence of tau condensates, 20 nM mScarlet-tau in assay buffer (20 mM HEPES pH 7.4, 40 mM KCl, 2 mM MgCl_2_ 1 mM EGTA, 10% (v/v) glycerol, 0.5% (w/v) pluronic F127, 2 mM ATP, 1 mM TCEP, 0.2 mg/mL casein, and 10 μM taxol) supplemented with an oxygen scavenger mix to remove free oxygen from solution (7.5 U/μL catalase, 0.15 U/μL glucose oxidase, 20 mM D-glucose) was perfused into the chamber with microtubules already immobilized on the surface and incubated for 10 min to allow the chamber to equilibrate. Subsequently, a mixture of 1 nM KIF5B-GFP and 20 nM mScarlet-tau was perfused into the chamber. For kinesin-1 motility assays on tau-free microtubules in the presence or absence of MAP7, KIF5B-GFP (or KIF5B-GFP and 1 nM MAP7-LD655) was diluted in assays buffer and perfused into the chamber with microtubules already immobilized on the surface. For kinesin-1 motility assays in the presence of tau condensates and MAP7, a mixture of 20 nM mScarlet-tau and 1 nM MAP7-LD655 in assay buffer was perfused into the chamber with microtubules already immobilized on the surface and incubated for 10 min to allow the chamber to equilibrate. Next, a mixture of 20 nM sfGFP-tau, 1 nM MAP7-LD655, and 1 nM K560-GFP was perfused into the chamber. Kinesin motility was recorded at a framerate of 5 or 8 Hz for 30-60s at 100 ms exposure per frame. For the experiments without tau, the same kinesin concentrations and imaging settings were used as those used in the experiments with tau.

tau-MAP7 intensity analysis was performed by extracting the intensity values from line scans along microtubules in FIJI ^96^. The tau and MAP7 intensity values per pixel were plotted against each other to generate relative intensity distributions. Data analysis of kinesin motility was performed as described in the previous section. To quantify run length and the percentage of processive KIF5B molecules moving though tau condensates in the presence of MAP7, a displacement cutoff of 0.3 μm was used. Therefore, Fig. 6e shows the percentage of molecules whose run length exceeds 0.3 μm.

### Tau single-molecule diffusion measurements

Flow chambers were prepared as described in the previous sections. tau condensates were first pre-formed on microtubules immobilized on the surface of the flow chamber as described above, either at 1 nM sfGFP-tau (island tau condensate regime) or 50 nM sfGFP-tau (complete microtubule coverage) in assay buffer assays buffer (20 mM HEPES pH 7.4, 75 mM KCl, 2 mM MgCl_2_ 1 mM EGTA, 10% (v/v) glycerol, 0.5% (w/v) pluronic F127, 2 mM ATP, 1 mM TCEP, 0.2 mg/mL casein, and 10 μM taxol) supplemented with an oxygen scavenger mix to remove free oxygen from solution (7.5 U/μL catalase, 0.15 U/μL glucose oxidase, 20 mM D-glucose). Next a mixture of either 10 pM mScarlet-tau and 1 nM sfGFP-tau (island tau condensate regime) or 200 pM mScarlet-tau and 100 nM sfGFP-tau (complete microtubule coverage) were perfused in the flow chamber. Single mScarlet-tau molecules were imaged continuously in the 561 nm channel at the frame rate of 10 Hz for 30 s with a 100 ms exposure per frame. Single tau molecules were tracked and analyzed in FIESTA. Mean squared displacement values were calculated using a built-in function in FIESTA. The values were plotted in GraphPad Prism 9 and fit to a linear one-dimensional diffusion equation: MSD = 2Dτ + C, where τ is a lag time, D is a diffusion constant, and C is an offset. Diffusion constants of individual tau molecules were obtained from FIESTA tracks using a custom-written MATLAB script (GitHub). Landing rate was quantified by dividing the number of molecules that interact with a microtubule by a product of the microtubule length, total measurement time, and the soluble tau concentration. Dwell times were extracted from the tracking data. Similar to run length quantification in the analysis of kinesin motility, only dwell times of molecules whose entire path can be observed during imaging were counted; i.e., molecules that landed on a microtubule before the beginning of imaging or dissociated after its end were not considered.

### MAP7 microtubule binding

Flow chambers were prepared as described in the previous sections. 20 nM sfGFP-tau in assays buffer (20 mM HEPES pH 7.4, 40 mM KCl, 2 mM MgCl_2_ 1 mM EGTA, 10% (v/v) glycerol, 0.5% (w/v) pluronic F127, 2 mM ATP, 1 mM TCEP, 0.2 mg/mL casein, and 10 μM taxol) supplemented with an oxygen scavenger mix to remove free oxygen from solution (7.5 U/μL catalase, 0.15 U/μL glucose oxidase, 20 mM D-glucose) was perfused into the chamber with microtubules already immobilized on the surface, and incubated for 10 min to allow the chamber to equilibrate. Then a mixture of 20 nM sfGFP-tau and 2.5 pM MAP7-LD655 in assay buffer was perfused into the chamber. Single MAP7-LD655 molecules were imaged continuously in the 647 nm channel at a framerate of 4.8 Hz for 5 min at 100 ms exposure per frame. Single MAP7 molecules were tracked and analyzed in FIESTA. The analysis was performed as described in the previous section.

### Image acquisition

All images were acquired on a Nikon Ti-E or Nikon Ti2 microscope equipped with a TI-TIRF adapter (Nikon) and EMCCD cameras (Andor iXon 897 or iXon 888). The excitation light was provided by a 488 nm Coherent CUBE for GFP operating at 200 mW, 561 nm Coherent CUBE for mScarlet operating at 300 mW, and 640 nm Coherent CUBE for HiLyte 647 or LD655 operating at 200 mW before being coupled into the TIRF module *via* an optical fiber. The excitation light was focused at the back focal plane of a 100X 1.49 NA oil objective (Nikon CFI Apo TIRF 100x). The excitation light was split from the emission by a quad band dichroic DI01/405/488/561/635 (Semrock) for imaging of sfGFP-tau, mScarlet-tau, or MAP7-LD655 or by a bandpass filter set ET-DsRed (ET545/30x/T570lp/ET620/60m, Chroma) for imaging of KIF1A-mScarlet. The emission was further split by an FF560-FDi02 (Semrock) filter for simultaneous imaging of sfGFP-tau and KIF1A-mScarlet or mScarlet-tau (488 and 561 channels) or FF640 filter (Semrock) for simultaneous imaging of mScarlet-tau and MAP7-LD655 (488 and 640 channels) and further filtered with an FF01-550/88 (Semrock) or 525/50 for the 488 channel, a 568R-25 for 568 channel and an FF01-642/LP (Semrock) or 647R for the 640 channel.

## AUTHOR CONTRIBUTIONS

I.Z. purified tau, tubulin, MAP7 and kinesins, performed and analyzed all assays. K.O.-M. provided plasmids, tau protein for initial experiments and MAP9. A.S. purified motors, tubulin and enzymes. A.R.-M. conceived, initiated, coordinated and supervised the project. I.Z. and A.R-M. interpreted data and wrote the manuscript. All authors read, edited the manuscript and approved final version.

## ACKNOWLEDGMENTS

We thank D.-Y. Lee from the Biochemistry Core (National Heart, Lung and Blood Institute, NHLBI) for help with mass spectrometry, J. Spector (National Institute of Neurological Disorders and Stroke, NINDS) for help with microscopes, A. Yildiz (University of California, Berkeley) for MAP7 and KIF5B plasmids, Adriaan Bax and Attila Szabo (National Institute of Diabetes and Digestive and Kidney diseases) for helpful discussions. A.R.-M. is supported by intramural programs of NINDS and NHLBI. The contributions of the NIH authors were made as part of their official duties as NIH federal employees, are in compliance with agency policy requirements, and are considered Works of the United States Government. However, the findings and conclusions presented in this paper are those of the authors and do not necessarily reflect the views of the NIH or the U.S. Department of Health and Human Services.

## DATA, CODE AND MATERIAL AVAILABILITY

Scripts used for analysis of tau diffusion and kinesin motility parameters were deposited on GitHub (https://github.com/RollmecakLab/Diffusion-and-track-analysis). All plasmids used in this study will be deposited to Addgene. Data will be archived and deposited as supplementary data files.

## Extended Data

Extended Data Figures 1-11

**Extended Data Fig. 1.**
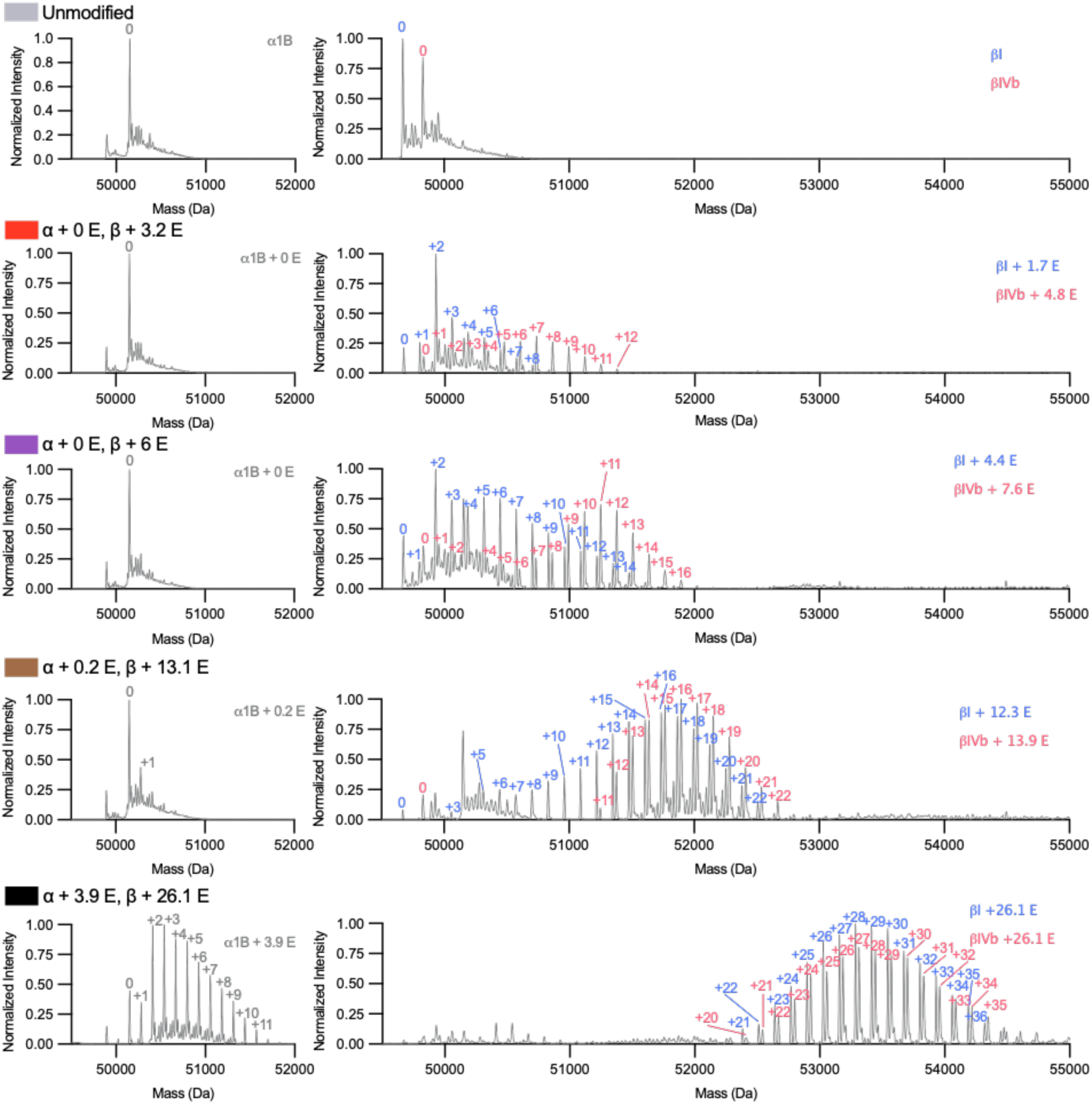
Mass spectrometry of differentially glutamylated microtubules. LC-MS of glutamylated microtubules used in assays. There are one major α (α1B) and two β (βI and βIVb) isotypes in these preparations. Throughout, numbers of glutamates added to α-tubulin are indicated in grey, and to βI and βIV-tubulin in blue and red, respectively. The weighted mean of the number of glutamates <n^E^> added to α-and β-tubulin (overall and separately for the two β-tubulin isoforms) are denoted α + <n^E^> and β+<n^E^> (Methods)

**Extended Data Fig. 2.**
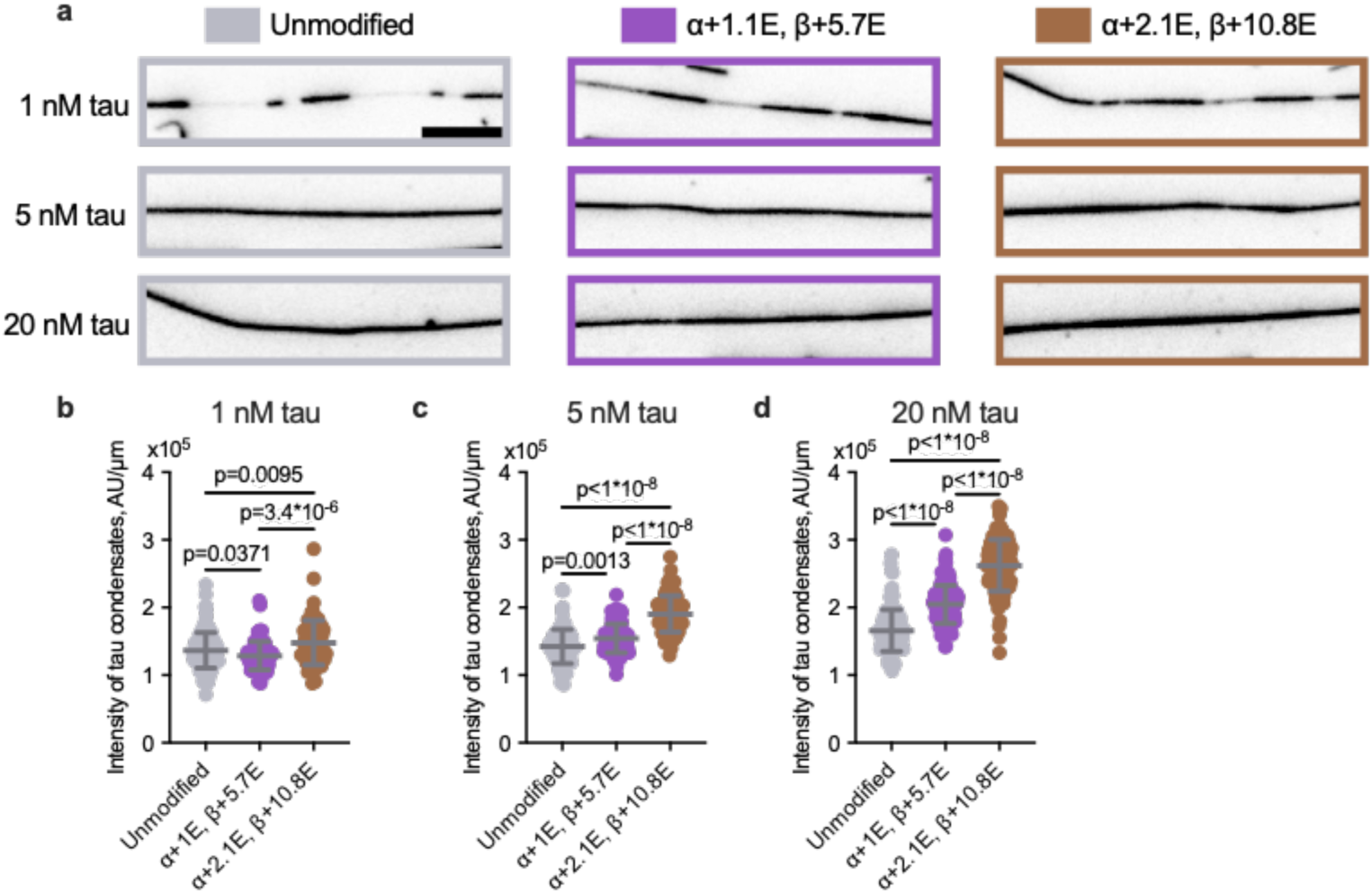
Tau condensate formation as a function of tubulin glutamylation. **a**, Condensate formation at increasing sfGFP-tau concentrations (top to bottom) and increasing glutamylation levels (left to right). **b**, Intensity of sfGFP-tau in condensates at 1 nM total tau concentration, n = 143, 115 and 72 microtubules for unmodified, <n_E_>_β_ ∼ 5.7 and 10.8, respectively. **c**, Intensity of sfGFP-tau in condensates at 5 nM total tau concentration, n = 119, 98 and 104 microtubules for unmodified, <n_E_>_β_ ∼ 5.7 and 10.8, respectively. **d**, Intensity of sfGFP-tau in condensates at 20 nM total tau concentration, n = 115, 123 and 114 microtubules for unmodified, <n_E_>_β_ ∼ 5.7 and 10.8, respectively; p-values determined by one-way ANOVA. 6 independent experiments for unmodified and 3 for the <n_E_>_β_ ∼ 5.7 and 10.8 conditions. Data points colored according to glutamylation levels, defined in (**a**).

**Extended Data Fig. 3.**
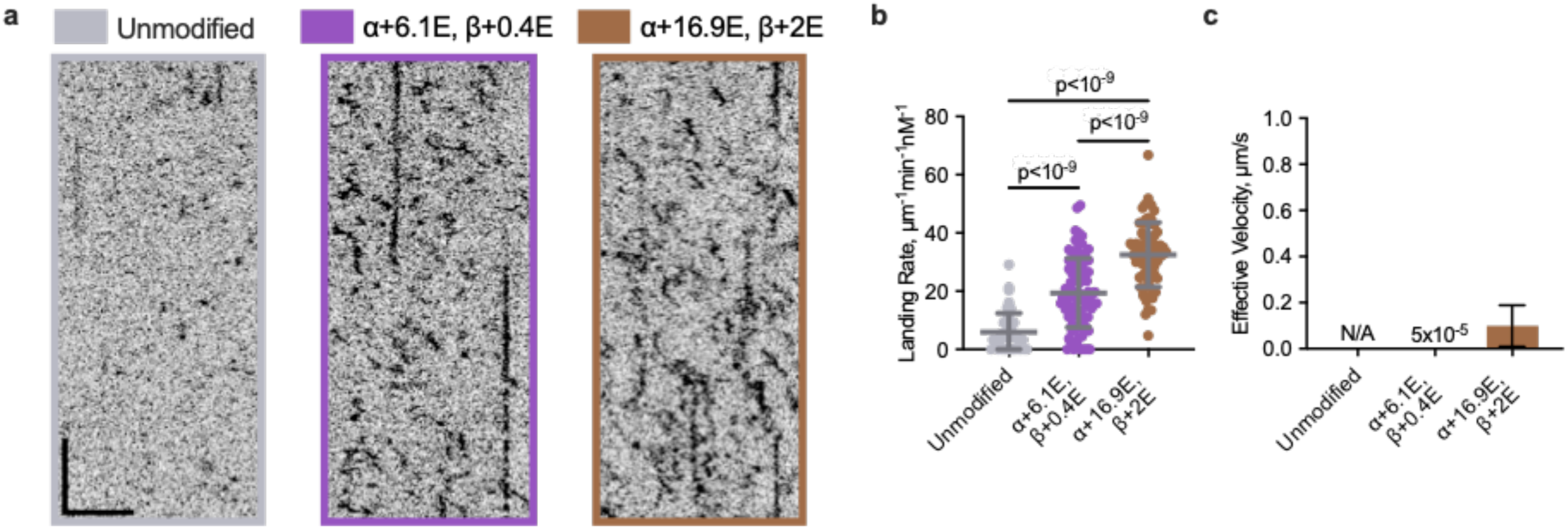
Glutamylation on α-tubulin stimulates KIF1A binding but does not enable processive motility through tau condensates. **a**, Kymographs of KIF1A-mScarlet motility on unmodified and glutamylated microtubules covered in tau condensates. 20 nM sfGFP-tau. Scale bars: horizontal, 5 μm; vertical, 5 s. **b**,**c**, KIF1A landing rate (**b**) and effective velocity (**c**) on unmodified and microtubules glutamylated on primarily α-tubulin, n = 49, 87, and 72 microtubules for unmodified, <n_E_>_α_ ∼ 6.1 and 16.9, respectively; p-values determined by one-way ANOVA (see Methods). 6 independent experiments for unmodified and 3 for <n_E_>_α_ ∼ 6.1, and 16.9 microtubule conditions. Data points colored according to glutamylation levels, defined in (**a**).

**Extended Data Fig. 4.**
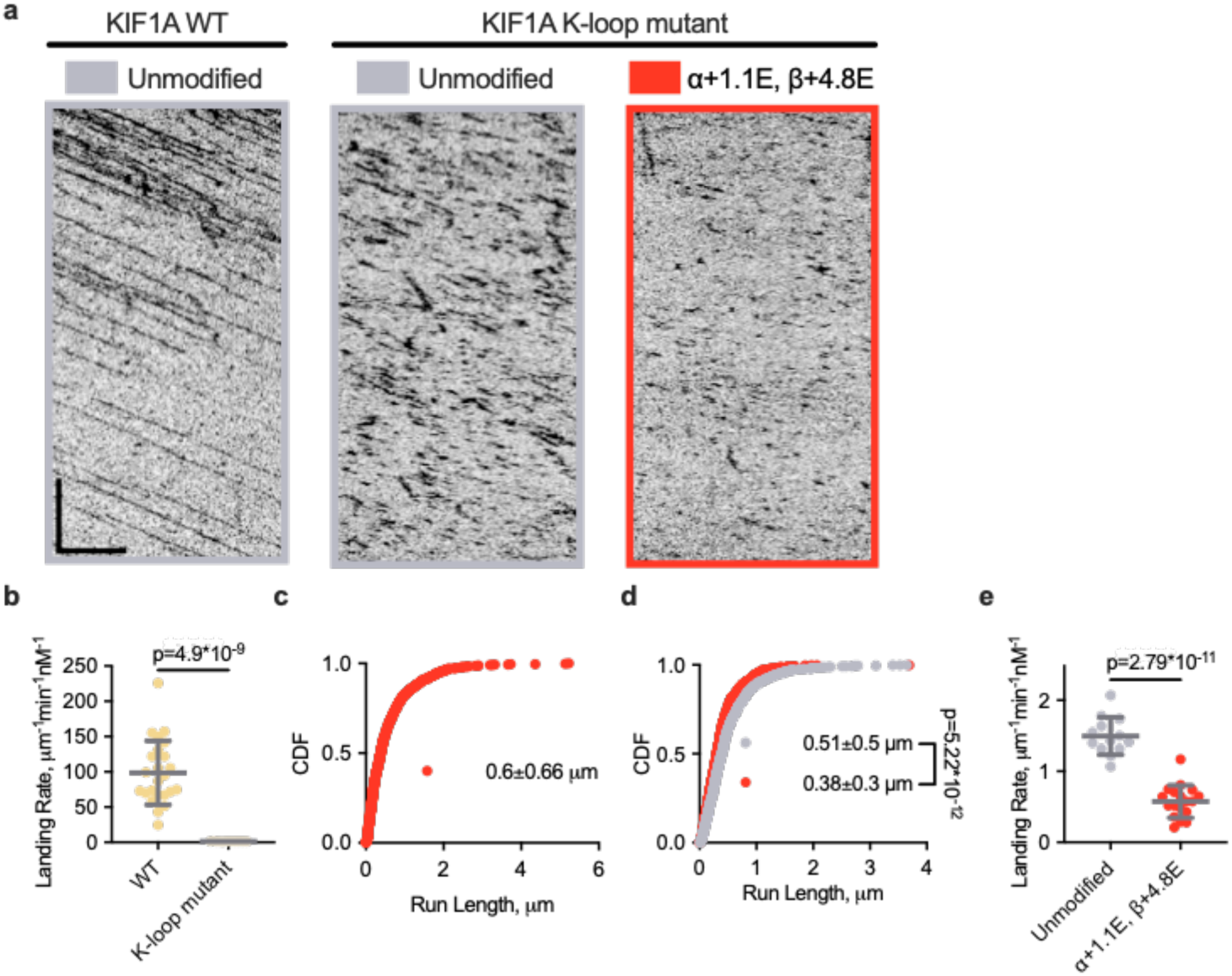
Substitution of the KIF1A K-loop with that from KIF5B impairs motility on glutamylated microtubules. **a**, Kymographs of wild-type KIF1A motility on unmodified tau-free microtubules (left) and KIF1A K-loop mutant on tau-free unmodified and glutamylated microtubules. Scale bars: horizontal, 5 μm; vertical, 5 s. **b**, Landing rate of WT and KIF1A K-loop mutant on unmodified microtubules; n = 24 and 13 microtubules for WT and KIF1A K-loop mutant, respectively. **c**, CDF of run length for wild-type KIF1A on glutamylated microtubules; n = 737 molecules for <n_E_>_β_ ∼ 4.8. Mean values for run lengths ± S.D. shown next to the graph. **d**, CDF of run length for the KIF1A K-loop mutant on unmodified and glutamylated microtubules; n = 1167 and 1012 molecules for unmodified and <n_E_>_β_ ∼ 4.8, respectively. Mean values for run lengths ± S.D. shown next to the graph; p-value determined by Kolmogorov-Smirnov test. **e**, Landing rate of the K-loop KIF1A mutant, n = 13 and 18 microtubules for unmodified and <n_E_>_β_ ∼ 4.8, respectively; p-value determined by two-tailed t-test, respectively; p-value determined by two-tailed t-test. 3 independent experiments for all conditions. Data points colored according to glutamylation levels, defined in (**a**).

**Extended Data Fig. 5.**
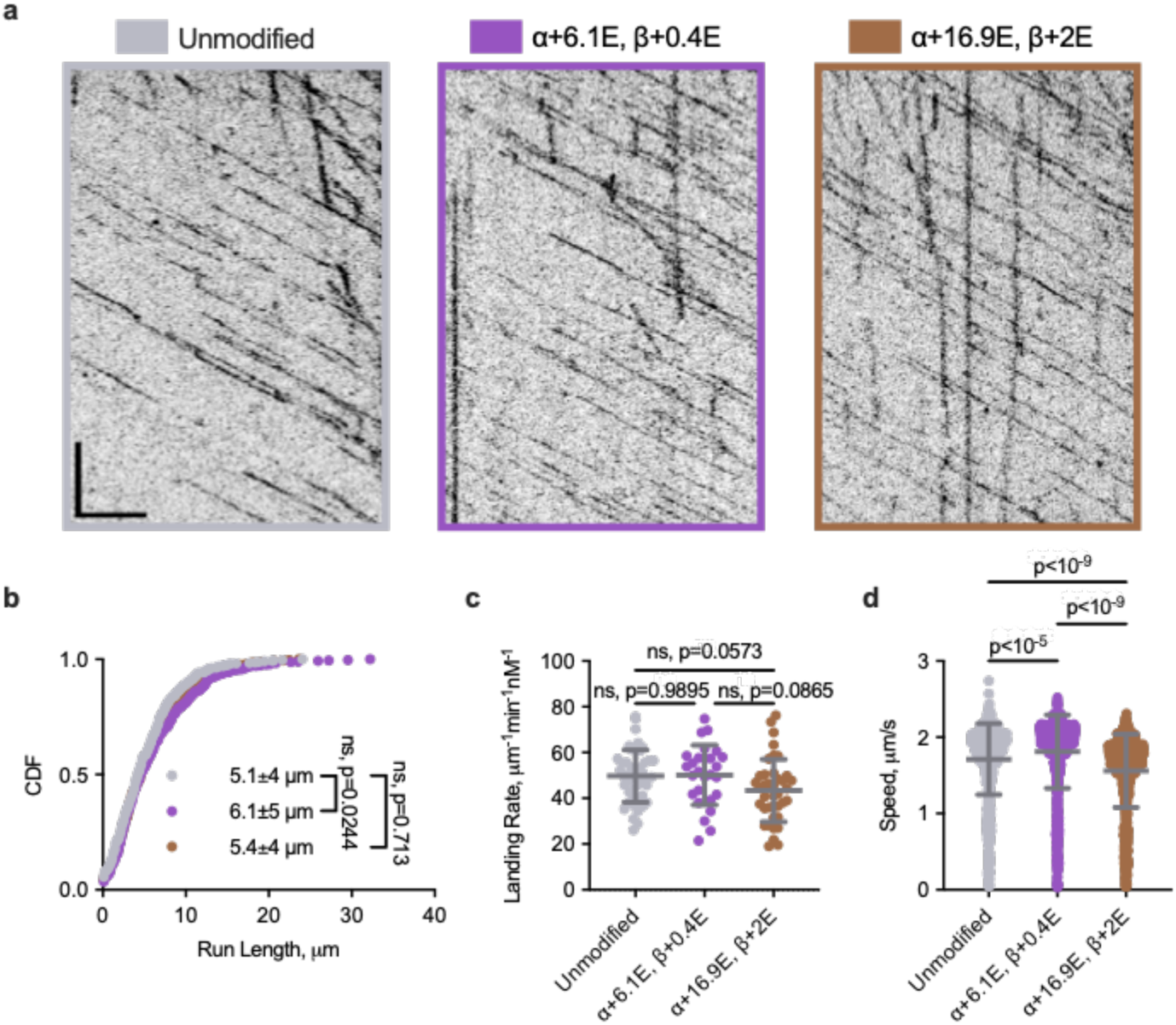
Glutamylation of α-tubulin does not stimulate KIF1A motility on tau-free microtubules. **a**, Kymographs of KIF1A-mScarlet motility at increasing α-tubulin glutamylation levels. Scale bars: horizontal, 5 μm; vertical, 5 s. **b**, CDF of KIF1A run length on differentially glutamylated microtubules, n = 450, 423, and 427 molecules for unmodified, <n_E_>_α_ ∼ 6.1, and 16.9, respectively; mean values ± S.D. shown next to the graph; p-values determined by Kolmogorov-Smirnov test. **c**, KIF1A landing rate, n = 47, 27, and 39 microtubules for unmodified, <n_E_>_α_ ∼ 6.1 and 16.9, respectively. **d**, KIF1A speed, n = 1235, 964, and 968 for unmodified, <n_E_>_α_ ∼ 6.1 and 16.9, respectively; p-values determined by one-way ANOVA; p-values determined by one-way ANOVA. 6 independent experiments for unmodified and 3 for <n_E_>_α_ ∼ 6 and 16.9 modified microtubule conditions. Data points colored according to glutamylation levels, defined in (**a**).

**Extended Data Fig. 6.**
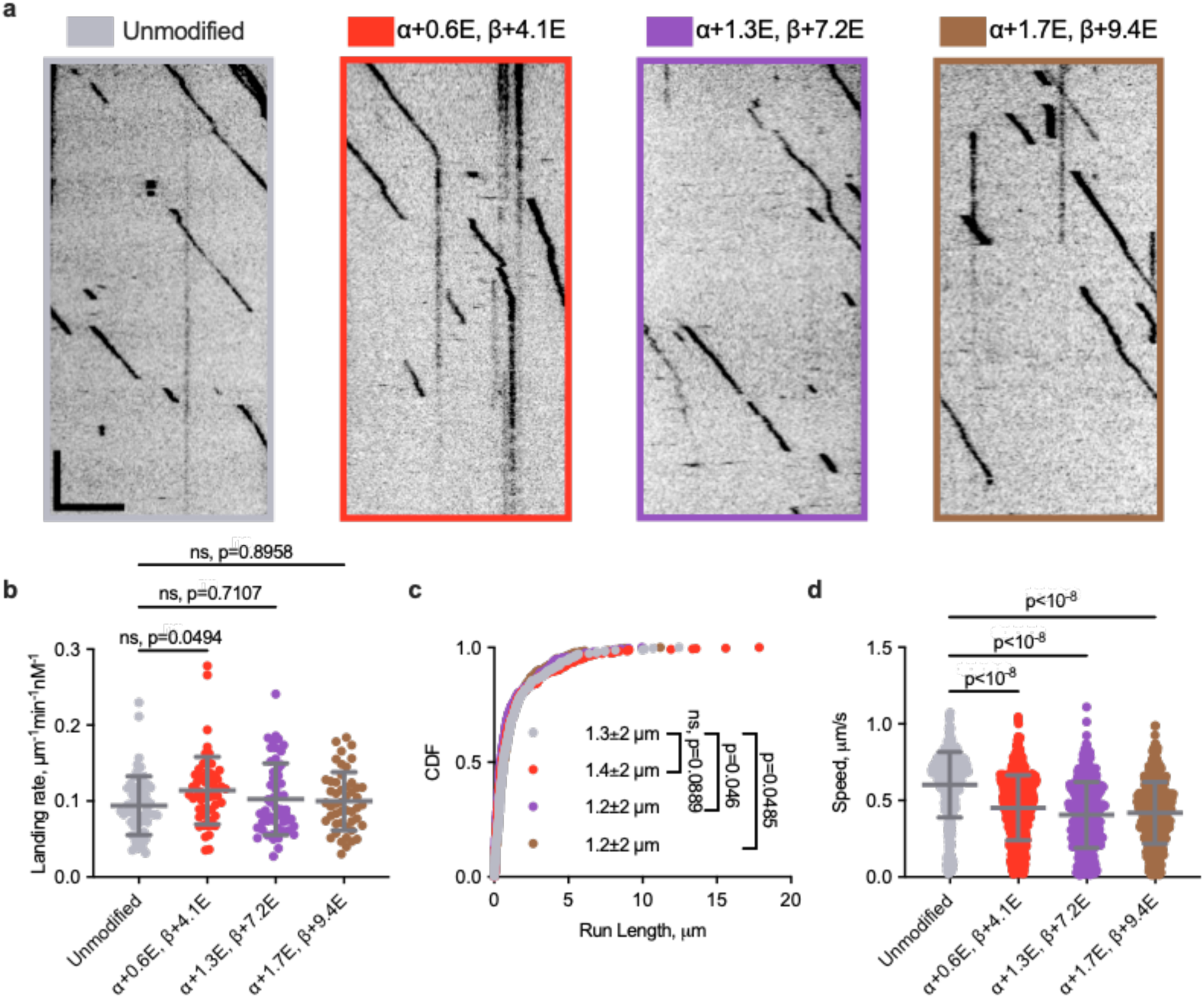
Glutamylation of β-tubulin has minimal effect on kinesin-1 KIF5B motility on tau-free microtubules. **a**, Kymographs of KIF5B-GFP motility on tau-free microtubules. Scale bars: horizontal, 5 μm; vertical, 5 s. **b**, Landing rate on tau-free microtubules, n = 62, 59, 55, and 49 microtubules for unmodified, <n_E_>_β_ ∼ 4.1, 7.2 and 9.4, respectively. **c**, CDF of KIF5B run length on tau-free microtubules; mean values ± S.D. shown next to the graph, p-values determined by Kolmogorov-Smirnov test. **d**, Speed on tau-free microtubules, n = 478, 666, 379 and 429 molecules for unmodified, <n_E_>_β_ ∼ 4.1, 7.2 and 9.4, respectively; p-values determined by one-way ANOVA. 3 independent experiments for all conditions. Data points colored according to glutamylation levels, defined in (**a**). See Supplementary Video S3.

**Extended Data Fig. 7.**
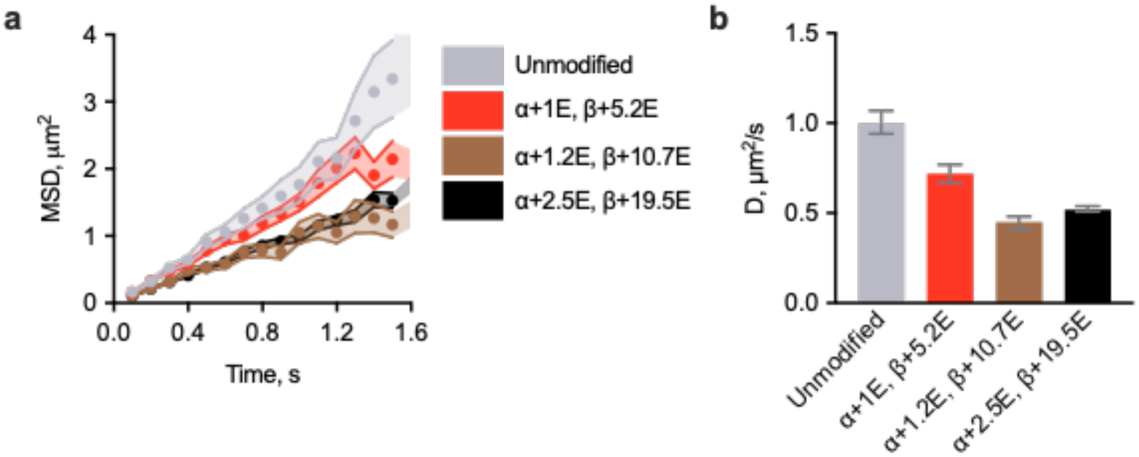
Effects of glutamylation on tau mobility outside condensates. **a**, MSD analysis of single mScarlet-tau molecules (10 pM) outside condensates at 1 nM total tau concentration, n = 90, 58, 42, and 90 molecules for unmodified (gray), <n_E_>_β_ ∼ 5.2 (red), 10.7 (magenta) and 19.5 (brown), respectively. Shaded areas show S.D. For details see Methods. **b**, Diffusion constant of tau outside condensates, extracted from the MSD fits in (**a**). 7 independent experiments for unmodified, 3 for <n_E_>_β_ ∼ 5.3 and 10.7, and 6 for <n_E_>_β_ ∼ 19.5 microtubule conditions. Data points colored according to glutamylation levels, defined in (**a**).

**Extended Data Fig. 8.**
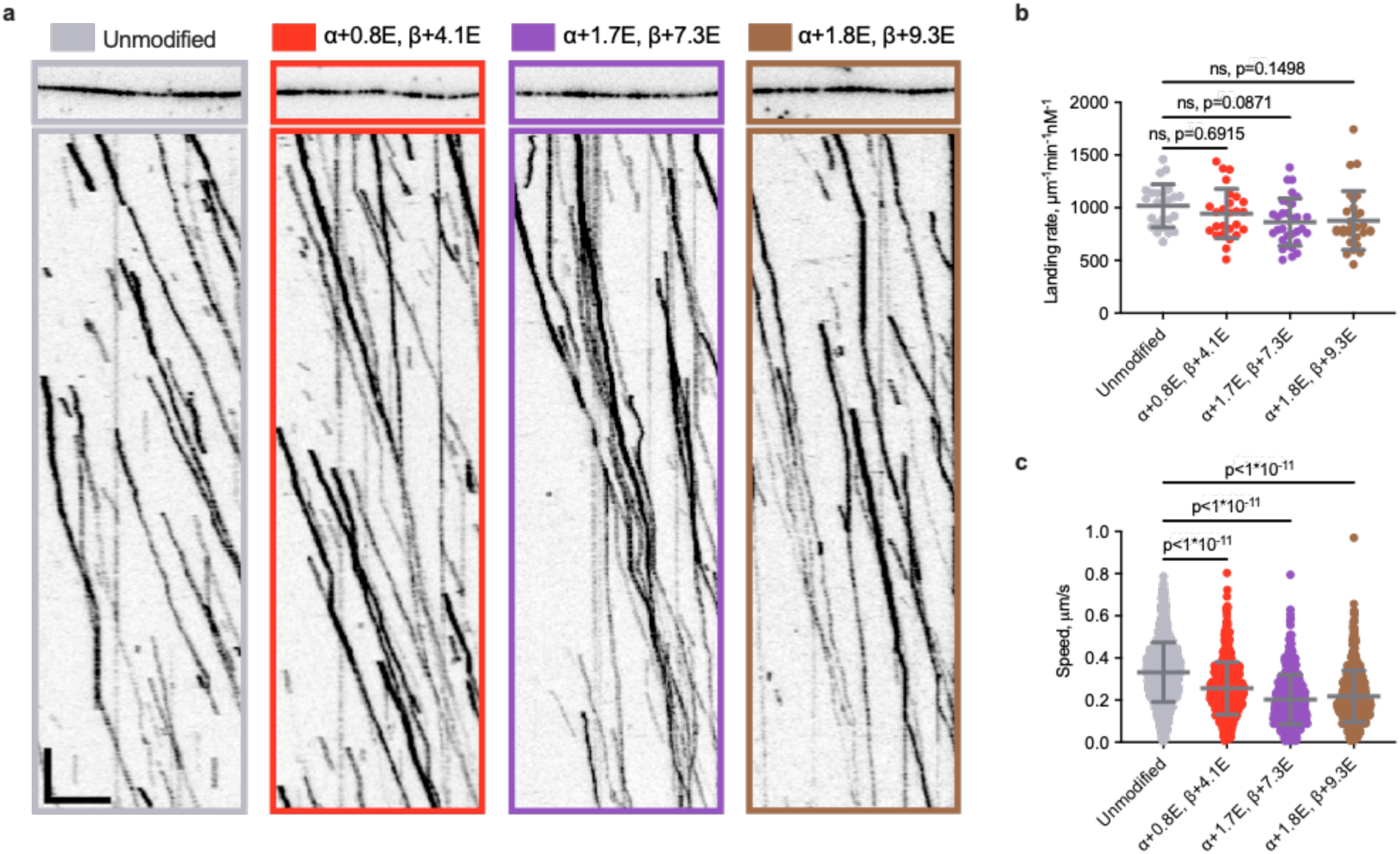
Glutamylation of β-tubulin has minimal effect on kinesin-1 KIF5B motility in the presence of MAP7 on tau-free microtubules. **a**, Kymographs of KIF5B-GFP motility on tau-free microtubules in the presence of 10 nM MAP7 labeled with LD655 (Methods) at different glutamylation levels. Scale bars: horizontal, 5 μm; vertical, 5 s. **b**, KIF5B landing rate in the presence of 10 nM MAP7 at increasing glutamylation levels, n = 25, 26, 31, and 29 microtubules for unmodified, <n_E_>_β_ ∼ 4.1, 7.3 and 9.3, respectively. **c**, KIF5B speed in the presence of 10 nM MAP7 at increasing glutamylation levels, n = 921, 604, 553, and 575 molecules for unmodified, <n_E_>_β_ ∼ 4.1, 7.3 and 9.3, respectively; p-values determined by one-way ANOVA. 3 independent experiments for all conditions. Data points colored according to glutamylation levels, defined in (**a**). See Supplementary Video 6.

**Extended Data Fig. 9.**
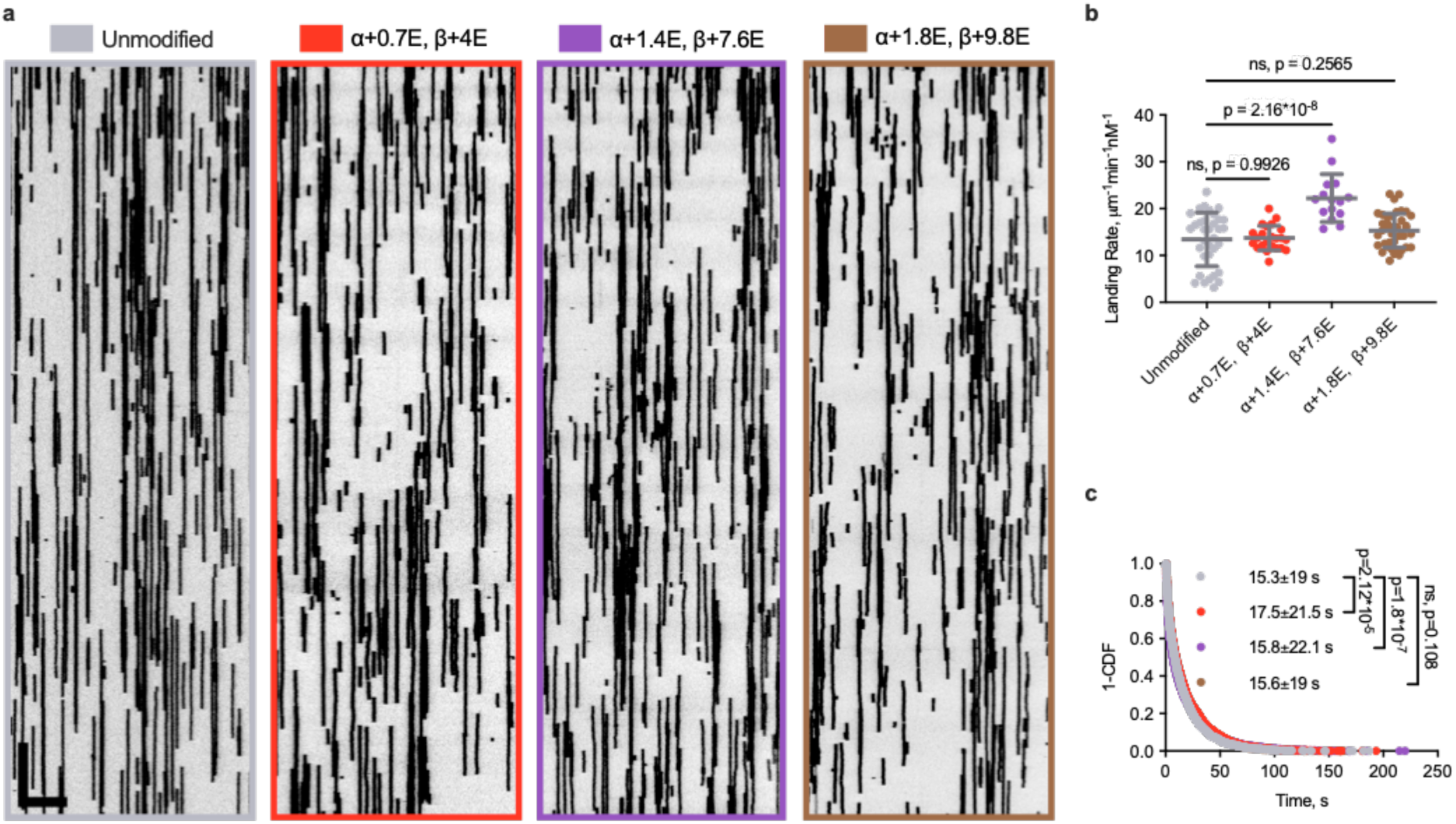
Glutamylation of β-tubulin does not affect MAP7 binding kinetics to microtubules covered in tau condensates. **a**, Kymographs of single LD655-labeled MAP7 molecules on differentially glutamylated microtubules covered in tau condensates; 20 nM sfGFP-tau concentration. **b**, MAP7 landing rate on differentially glutamylated microtubules covered in tau condensates; p-values determined by one-way ANOVA. **c**, Inverted CDF of dwell times of single MAP7 molecules on differentially glutamylated microtubules covered in tau condensates; mean values ± S.D. shown next to the graph; n = 4850, 5875, 5512, and 5772 molecules for unmodified, <n_E_>_β_ ∼ 4, 7.6 and 9.8, respectively; p-values determined by Kolmogorov-Smirnov test. 3 independent experiments for all conditions. Data points colored according to glutamylation levels, defined in (**a**). See Supplementary Video 7.

**Extended Data Fig. 10.**
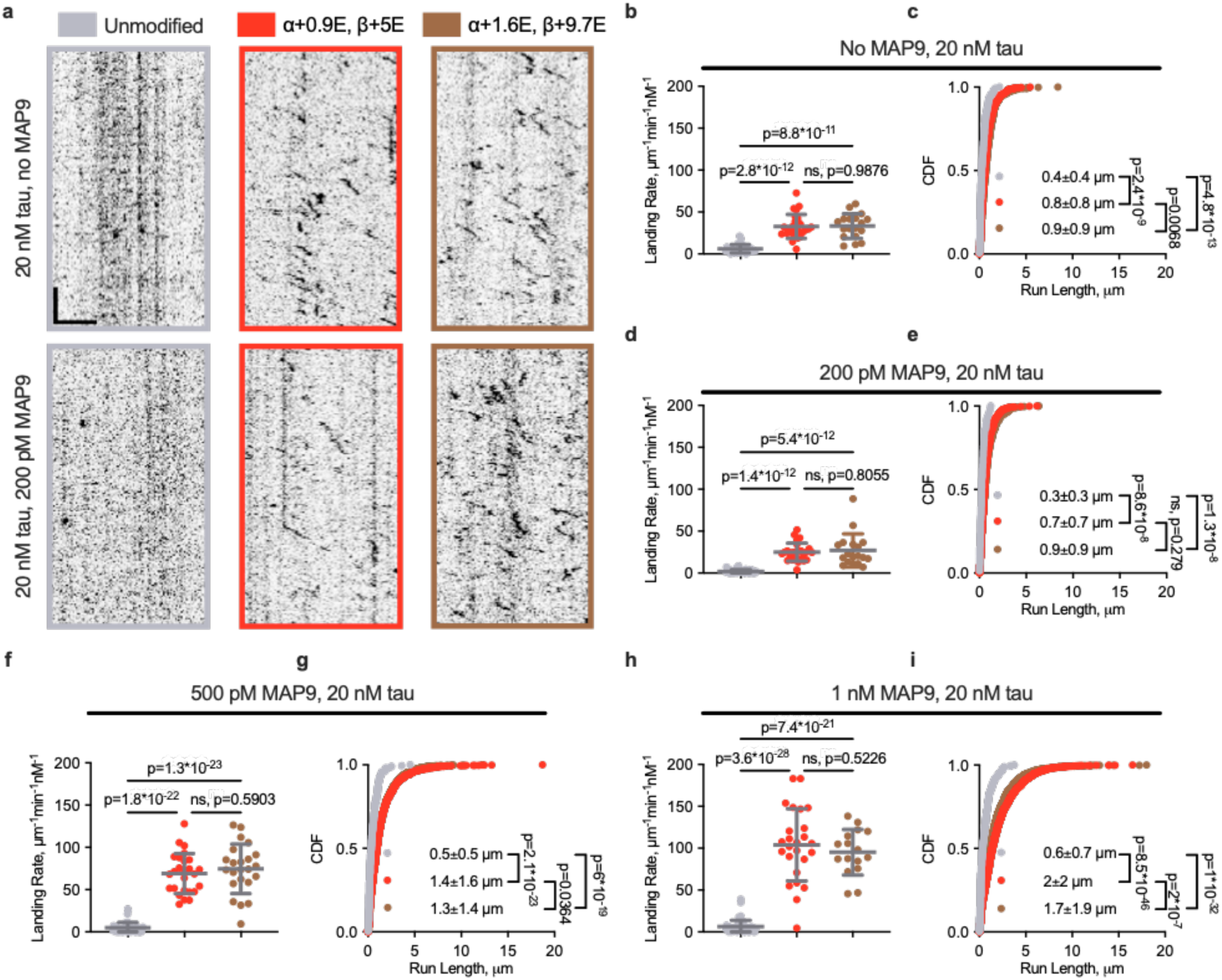
Glutamylation and MAP9 synergize to enable KIF1A motility on microtubules covered in tau condensates. **a**, Kymographs of KIF1A-mScarlet motility on differentially glutamylated microtubules covered in tau condensates in the absence of MAP9 (top) and presence of 0.2 nM MAP9; 20 nM sfGFP-tau concentration. Scale bars: horizontal, 5 μm; vertical, 5 s. **b**, KIF1A landing rate on differentially glutamylated microtubules covered in tau condensates and no MAP9; n = 30, 27, and 18 microtubules for unmodified, <n_E_>_β_ ∼ 5 and 9.7, respectively. **c**, CDF of KIF1A run length on differentially glutamylated microtubules covered in tau condensates and no MAP9; n = 169, 1022, and 852 molecules for unmodified, <n_E_>_β_ ∼ 5 and 9.7, respectively. **d**, KIF1A landing rate on differentially glutamylated microtubules covered in tau condensates in the presence of 0.2 nM MAP9; n = 43, 26, and 18 microtubules for unmodified, <n_E_>_β_ ∼ 5 and 9.7, respectively. **e**, CDF of KIF1A run length on differentially glutamylated microtubules covered in tau condensates in the presence of 0.2 nM MAP9; mean values ± SD shown next to the graph; n = 74, 761, and 549 molecules for unmodified, <n_E_>_β_ ∼ 5 and 9.7, respectively. **f**, KIF1A landing rate on differentially glutamylated microtubules covered in tau condensates in the presence of 0.2 nM MAP9; n = 44, 26, and 23 microtubules for unmodified, <n_E_>_β_ ∼ 5 and 9.7, respectively. **g**, CDF of KIF1A run length on differentially glutamylated microtubules covered in tau condensates in the presence of 0.5 nM MAP9; n = 226, 2085, and 2111 molecules for unmodified, <n_E_>_β_ ∼ 5 and 9.7, respectively. **h**, KIF1A landing rate on differentially glutamylated microtubules covered in tau condensates in the presence of 1 nM MAP9; n = 54, 25, and 16 microtubules for unmodified, <n_E_>_β_ ∼ 5 and 9.7, respectively. **i**, CDF of KIF1A run length on differentially glutamylated microtubules covered in tau condensates in the presence of 1 nM MAP9; mean values ± SD shown next to the graph; n = 307, 2639, and 2367 molecules for unmodified, <n_E_>_β_ ∼ 5 and 9.7, respectively. Scale bars: horizontal, 5 μm; vertical, 5 s; p-values determined by one-way ANOVA test in (**b**), (**d**), (**f**), and (**h**) and by Kolmogorov-Smirnov test in (**c**), (**e**), (**g**), and (**i**). 6 independent experiments for unmodified and 3 for <n_E_>_β_ ∼ 5 and 9.7 conditions. Data points colored according to glutamylation levels, defined in (**a**). See Supplementary Videos 8 and 9.

**Extended Data Fig. 11.**
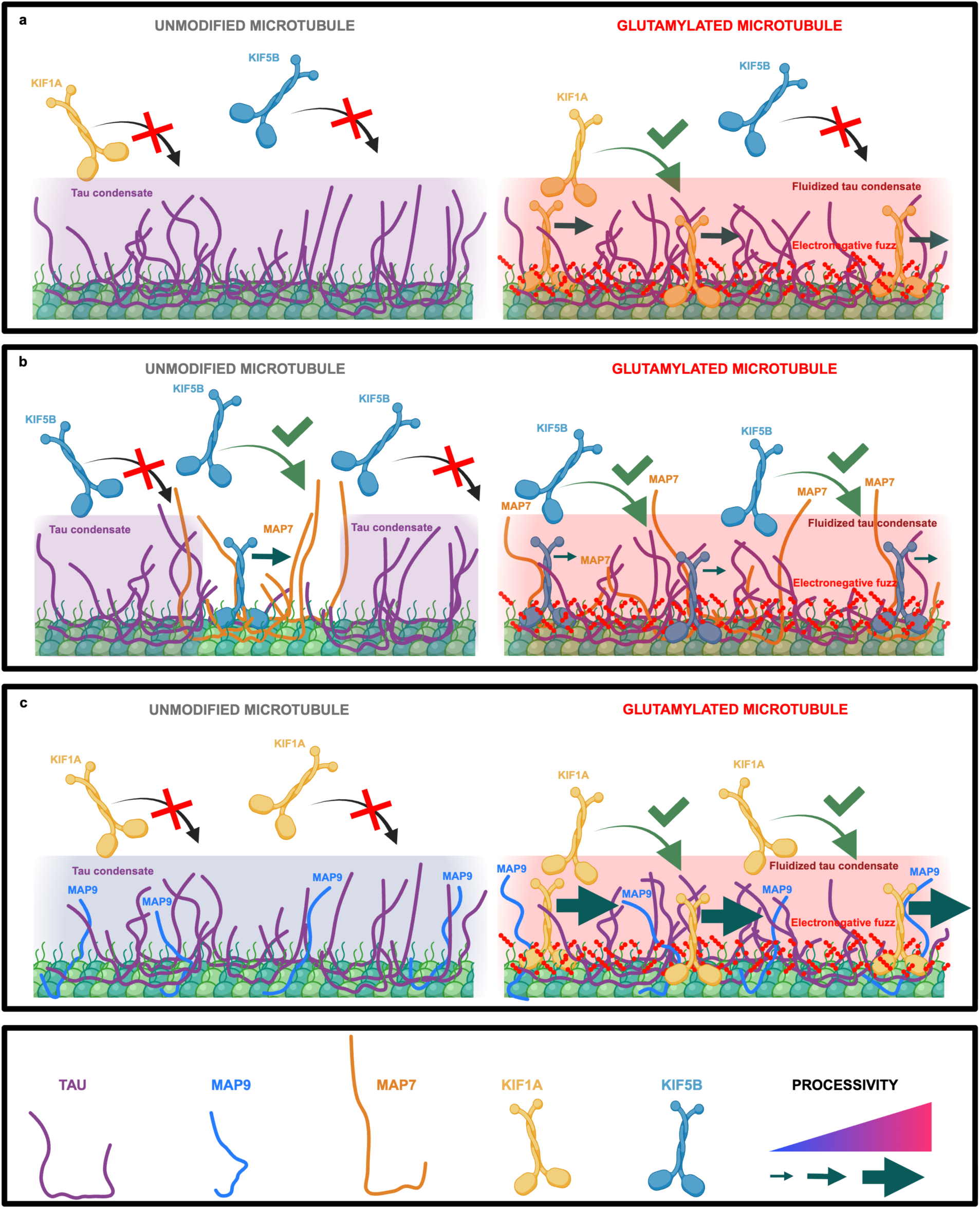
Glutamylation enables selective kinesin motility through tau barriers. **a**, Differential effects of glutamylation on kinesin-1 KIF5B and kinesin-3 KIF1A motility through tau condensates. **b**, MAP7 enables kinesin-1 KIF5B recruitment to tau condensates and supports limited motility. **c**, MAP9 synergizes with glutamylation to promote high recruitment and processivity of kinesin-3 KIF1A.

